# A modular and reusable model of epithelial transport in the proximal convoluted tubule

**DOI:** 10.1101/2021.11.11.468301

**Authors:** Leyla Noroozbabaee, Pablo J. Blanco, Soroush Safaei, David P. Nickerson

## Abstract

We review a collection of published renal epithelial transport models, from which we build a consistent and reusable mathematical model able to reproduce many observations and predictions from the literature. The flexible modular model we present here can be adapted to specific configurations of epithelial transport, and in this work we focus on transport in the proximal convoluted tubule of the renal nephron.

Our mathematical model of the epithelial proximal convoluted tubule describes the cellular and subcellular mechanisms of the transporters, intracellular buffering, solute fluxes, and other processes. We provide free and open access to the Python implementation to ensure our multiscale proximal tubule model is accessible; enabling the reader to explore the model through setting their own simulations, reproducibility tests, and sensitivity analyses.

## Introduction

Kidneys are vital organs and play an essential role in the overall homeostasis of the body in mammals. A human kidney is typically composed of one million nephrons, which are the primary functional unit of the kidney. The nephron consists of the renal corpuscle and the renal tubule. A renal tubule is a tubular structure composed of a single layer of epithelial cells divided into various functional segments. Each segment of the nephron has specific functions in the regulation of blood and urine composition. The proximal convoluted tubule (PCT) is considered one of the most significant functional segments in the nephron and a key contributor to pathologies such as hypertension and diabetes.

To gain a deeper insight into the mechanisms and to investigate any hypothesis regarding the underlying physiopathological conditions, such as hypertension, diabetes, or other kidney diseases, a virtual nephron model is an invaluable tool. A model like this, should subserve as a virtual laboratory, it should be inexpensive to run and should target the minimisation of animal experiments.

Many models of epithelial transport have been published which would be relevant and useful to integrate into a virtual nephron model, but often there is insufficient information in the literature to enable readers to reproduce the published observations and predictions, thus making their reuse in novel studies time consuming and resource intensive. Furthermore, as models evolve over time, any given version of a model is usually designed to investigate specific hypotheses. The various instances of a given model, or a family of models, are therefore inconsistent and require readers to search the literature to discover or infer the modifications required to integrate the models and successfully reproduce published results. Such modifications, for example, may be as trivial as changes in parameter values or alterations in of physical units, or as complex as specific assumptions made or mathematical formulations chosen. These problems hamper efforts to develop a virtual nephron model.

To help address this problem, we introduce here an integrative mathematical model of the PCT which reproduces the capabilities of existing renal models. We used a modular approach to build our model, which we believe will improve reusability while ensuring the segment function predicted by the source models can be reproduced. At the same time, this approach is sufficiently flexible and configurable to be able to adapt to different segments or epithelium. This model has been implemented in Python and is freely available under an open-source and permissive license at https://github.com/iNephron/W-PCT-E.

Alan Weinstein has provided a valuable resource for mechanistic modelling of many aspects of renal function through many publications [1–5, for example]. Our long-term goal is to make this resource available in a more reproducible and reusable form via the CellML standard [6,7] and with bond graph approaches [8–12] that ensure energy conservation as well as mass and charge conservation across different physical systems (biochemical, electrical, mechanical and metabolic). However, to bring Weinstein’s work into a consistent and unified mathematical framework and to understand the various parameter combinations and manipulations that have been used in a series of publications spanning 30+ years, we first implement his model in a Python scripting environment. This environment is described here to make this resource available to the renal modelling community.

## Design and Implementation

In this work, we follow the comprehensive PCT epithelial model collectively presented in Weinstein et al. [1–5] which we refer to here as the W-PCT-E. The W-PCT-E consists of four different stages. The first stage begins with the system’s geometrical definitions and equations of the selected components such as cell volume, lateral intercellular volume, basement membrane area, and epithelial thickness. These parameters are variable while the rest of the geometrical components, such as membrane area for the other regions, are constant (see Section vii). The second stage indicates five different intra-epithelial fluxes: water fluxes, convective fluxes, passive fluxes, coupled solute fluxes, and active (metabolically dependent) fluxes. The output from the first stage will be the input to the second stage, plus the specification of the membrane types and solutes that appear in the chemical reactions. The outcome of the second stage will be total membrane solute fluxes and the membrane water fluxes.

In the third stage, the W-PCT-E model establishes the system mass conservation through the differential equations within each compartment (or each membrane) with the total solute fluxes and water fluxes as the input for each membrane.

In the final stage, the W-PCT-E model updates the differential equations by applying more constrains by defining the buffer pairs, pH equilibrium, and electroneutrality of the system.

### I. Model Illustrations

Here, we define the various components in the modelling framework in the W-PCT-E system, including all equations, definitions, and assorted tables of constants and variables that appear in the epithelial model’s compartments. As well, we introduce the ingredients present in our Python code and, whenever possible, the provenance of the parameter values. The comprehensive W-PCT-E model consists of the cellular and lateral intercellular compartments between luminal and peritubular solutions. Figure 2 displays a schematic representation of PCT epithelium and features both configurations, in which cellular and lateral intercellular (LIS) compartments line the tubule lumen. Within each compartment, the concentration of species (i) is designated *C*_*α*_(*i*), where *α* is lumen (M), lateral interspace (E), cell (I), or basal solution (S). The separating membranes are the combination of letters such that luminal cell membrane (lumen-cell membrane, MI), tight junction (ME), cell-lateral membrane (IE), interspace basement membrane (ES), or cell-basal membrane (IS). The order of the two letters indicates the positive direction of the mass flow. *J*_*αβ*_ and *J*_*ναβ*_ represent the solute flux and water flux, respectively, through the corresponding membrane; A is the corresponding membrane surface area; V is the volume; E is the trans-epithelial potential difference. Symbols are defined in the following sections as they are introduced by Weinstein et al. [4] in the epithelial PCT model. Intra-epithelial fluxes are designated *J*_*αβ*_(*i*), where *αβ* refers to the different membranes. Models can include many different solutes in various compartments. In this paper, according to [4], the model consists of 15 solutes, namely, Na^+^, K^+^, Cl^−^, HCO_3_ ^−^, CO_2_, H_2_CO_3_, HPO_4_^2 –^, H_2_PO_4_ ^−^, Urea, NH_3_, NH_4_^+^, H^+^, HCO_2_ ^−^, H_2_CO_2_, and glucose, as well as two impermeant species within the system; a nonreactive anion and a cytosolic buffer. The solutes considered in a specific simulation experiment can vary, leading to the dynamics of some of the solutes being ignored under certain conditions. There are 14 transporters (symporters, antiporters, complex transporters, and ATPases) that produce electrochemical fluxes in the W-PCT-E model.

**Figure 1.**
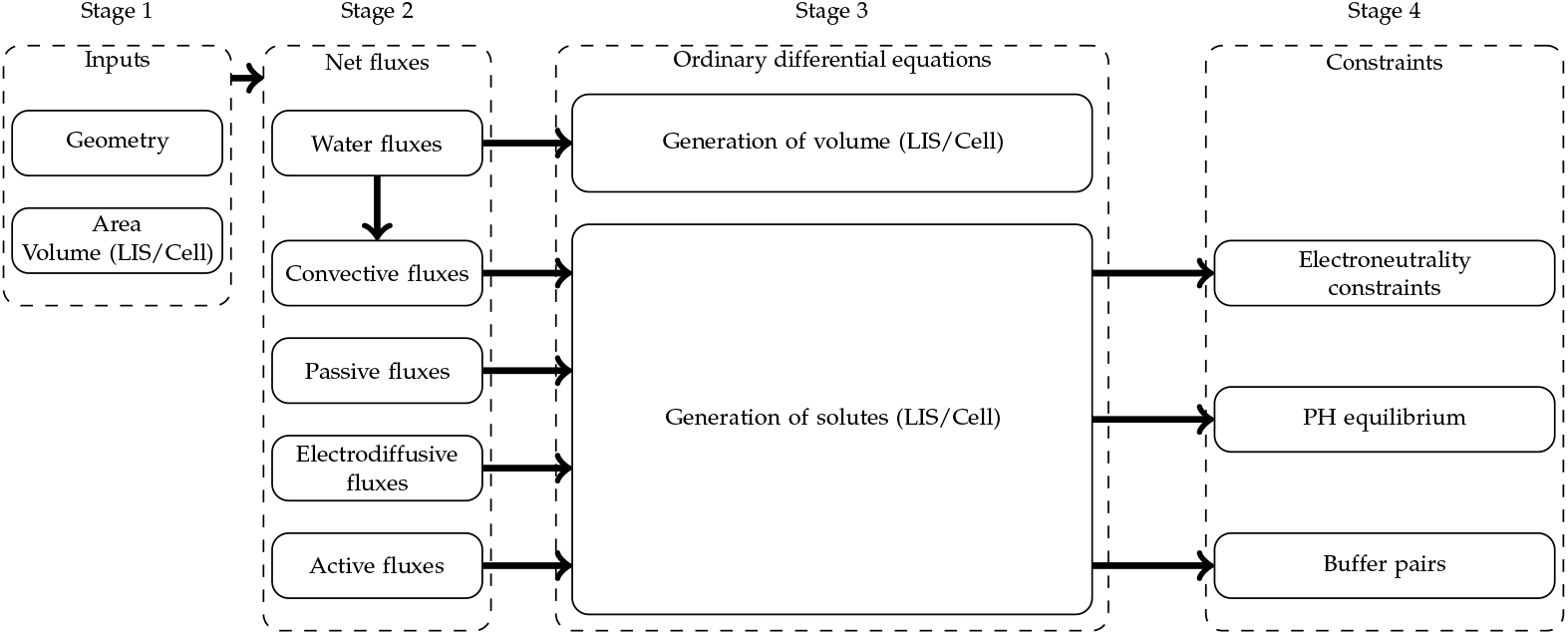
Block diagram of the epithelial model. The model begins with the geometrical inputs (see Section vii). The second box indicates the intra-epithelial fluxes; these fluxes will be involved in the mass conservation equations. The third box indicates the total mass conservation differential equations present in the epithelial model. The differential equations are, at last, be updated by the buffer pairs, pH equilibrium and electroneutrality of the system.

**Figure 2.**
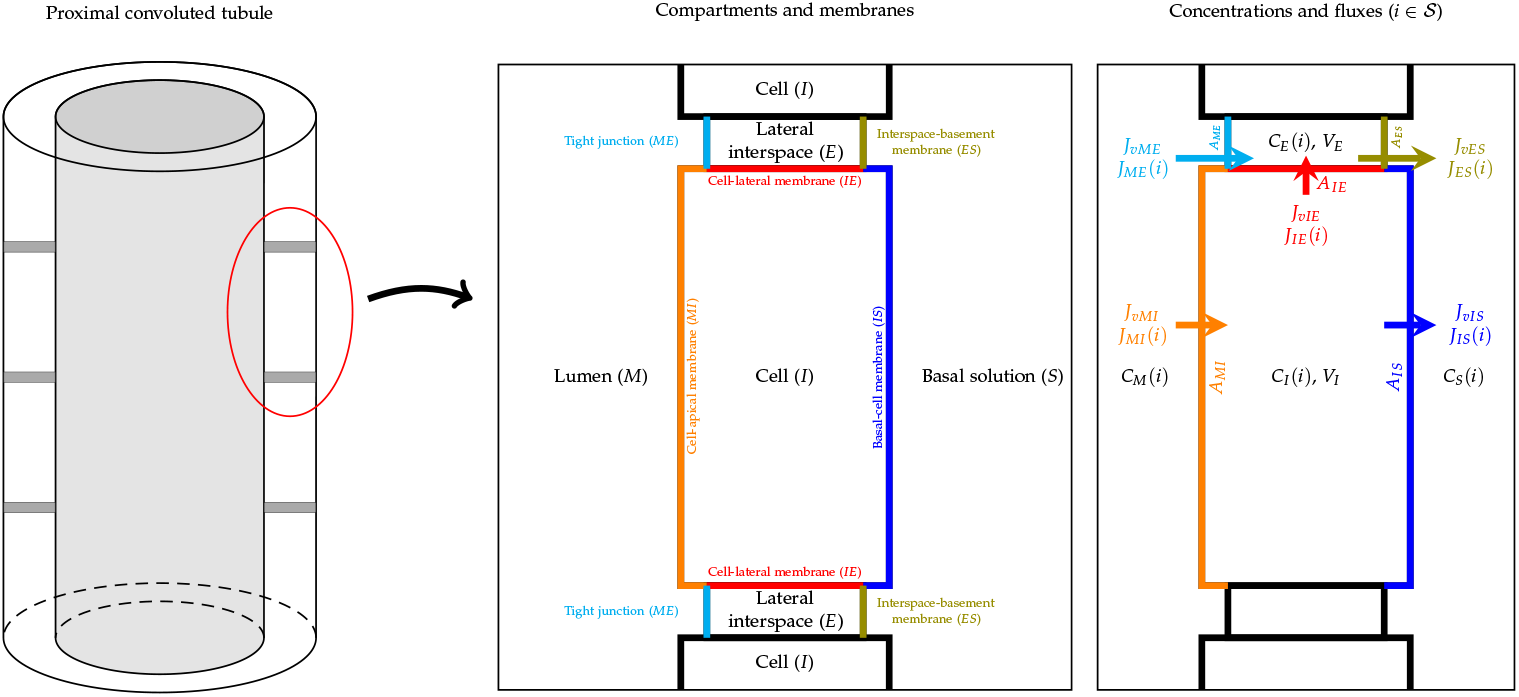
Schematic representation of proximal convoluted tubule (PCT) epithelium, consisting of cell and lateral intercellular space, and a tubule model, in the way lumen is lined by epithelium. Intra-epithelial fluxes are designated *J*_*αβ*_(*i*), where the subscript *αβ* refers to the different membranes and (*i*) refers to a specific solute.

Overall, there are six symporters on different membranes; two on the lumen-cell (MI): Sodium-Glucose (SGLT) and Sodium-Phosphate (NPT1), two on the cell-basal (IS): Potassium-Chloride (KCC4) and Sodium-Bicarbonate (NBC1), and two on the cell-lateral (IE): Potassium-Chloride (KCC4) and Sodium-Bicarbonate (NBC1). It is important to mention that cell-basal and cell-lateral membranes share a similar layout.

There are four antiporters on the lumen-cell membrane: Chloride/Bicarbonate (AE1), Chloride/Formate (Pendrin), Sodium/Hydrogen, and Sodium/Ammonium (NHE3). There are two complex exchangers on the cell-lateral membrane: Sodium-Bicarbonate/Chloride (NCBE) and one on the cell-basal membrane: Sodium-Bicarbonate/Chloride (NCBE). The Na/K pumps (NaK-ATPase) are located in both cell-basal and cell-lateral membranes. The H-pumps (H-ATPase) are located in the lumen-cell membrane, regulating the pH in the W-PCT-E model. We noticed two different approaches to translate the behaviour of Sodium/Hydrogen antiporter in Weinstein’s work. In the first approach, Weinstein et al. used the equivalent definition of two antiporters: Sodium/Hydrogen and Sodium/Ammonium [3]. In the second approach, reported in [4], the authors used a detailed model of the Na^+^/H^+^ antiporter from [13].

#### i. State Equations

First, we introduce the sets of compartments (𝒞) and membranes (ℳ) indexes

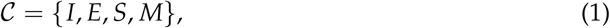

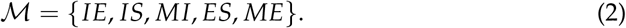

Indexes in ℳ might appear interchanged in some flux definitions, which means that the flux sign must be inverted. Now, we introduce the following set of solutes

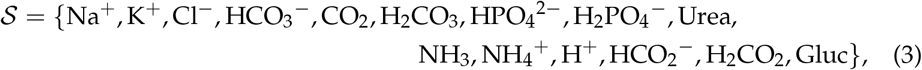

which is divided into non-reacting (*NR*) and reacting (*R*) solutes

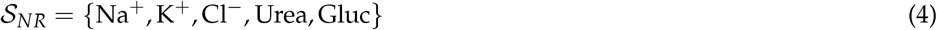

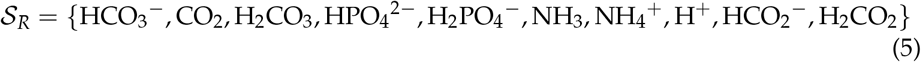

In the W-PCT-E model, the system of state equations represents two different compartments: cell and lateral intercellular spaces. To formulate the mass conservation equations within each compartment, the net generation of each species *S*_*α*_(*i*) is defined as an intermediate variable within the compartment [3]. The generation of **multiple reacting** solutes is the sum of the net exchange of the flux plus the accumulation of that solute in each compartment, that is

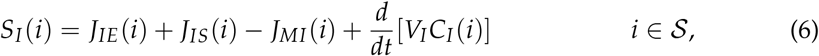

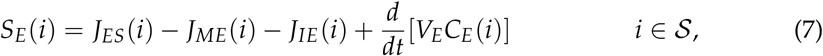

where *S*_*I*_ (*i*) and *S*_*E*_(*i*) indicate the generation for solute *i ∈ 𝒮* in the cell space and lateral interspace compartments, respectively. Within the epithelium, the flux of solute *i* across the membrane *αβ* is denoted as *J*_*αβ*_(*i*) [mmol. s^−1^. cm^−2^], *αβ ∈ ℳ* and *V*_*α*_ is the compartment volume per unit surface area [cm^3^/cm^2^. epithelium], *α*. The principle of mass conservation relating water fluxes and volume change for different compartments reads

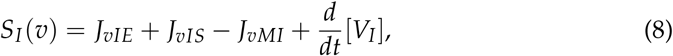

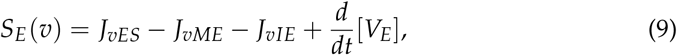

where *J*_*vαβ*_ [ml. s^−1^. cm^−2^] is denoted as the transmembrane volume flux. It is important to mention that for nonreacting solutes we have

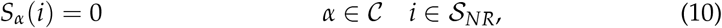

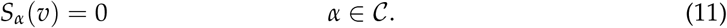

The mass conservation then defines the change of the concentration of the *i* species in the intracellular solution as the transport of solute *i* into and out of the cell through the apical and basolateral membrane. This is a direct transport of solutes through the membrane, and a contribution from convective transport due to the flow of water through the membranes. For such a nonreacting solute, the combination of (6) and (8) yields

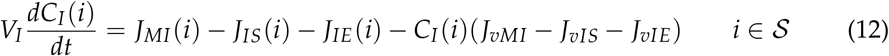

This equation holds for each solute *i ∈ 𝒮* being considered in a2 particular instantiation of the model. If required for a particular model, similar equations can be introduced for the solute concentrations in the mucosal and/or serosal solutions.

Consequently, from (8), the conservation of cellular water yields the equation below, with the rate of change of cell volume, *V*_*I*_, defined as

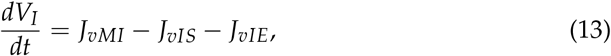

where each of the total membrane water fluxes, *J*_*vαβ*_, *αβ ∈ 𝒮*, is scaled (multiplied) by its respective membrane area to take into account the averaged behaviour of the representative membrane. For more detailed information of all various fluxes across the membrane, see Section II.

To calculate the concentration of the cellular buffer pairs, the following expression is employed [3]

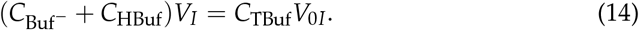

where *C*_Buf_^−^ and *C*_HBuf_ are cell buffer and protonated cell buffer concentration, respectively. *V*_*I*_ and *V*_0*I*_ indicate the cell volume and cell volume reference, respectively. To add more constraints on the proton cellular concentration, and to generate the paired equation, we have the following equilibrium equations

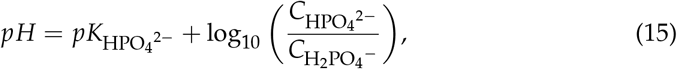

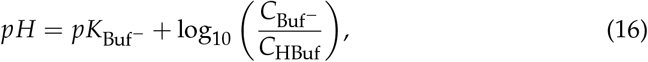

where 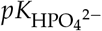 and 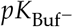 are the equilibrium constants for the phosphate and cell buffer pairs. Considering that the concentration of the H^+^ remains constant for all buffer pairs in the model, expressions (15) and (16) are combined to give

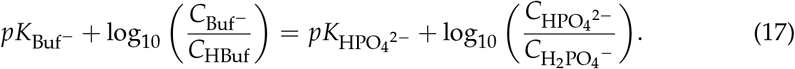

To include the cellular proton concentration in the proton mass conservation, *S*_*I*_ (H^+^), the following modification is applied to the equation (6)

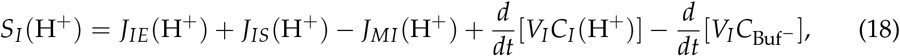

which can equivalently be written in the following form

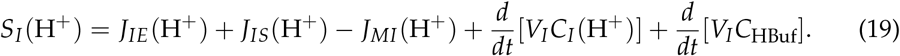

#### ii. Buffer Pairs and pH Equilibrium

The W-PCT-E model defines different types of buffer pairs, the mass conservation principle for the phosphate and formate buffer pairs takes the form

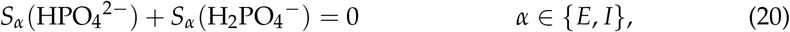

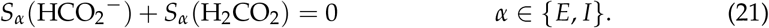

Similar equations apply to the ammonia pair within the lateral interspace and cell [3], that is *S*_*α*_(NH_3_) +

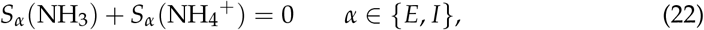

For an impermeant cytosolic buffer, Buf^−^, it is

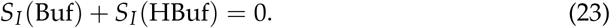

All buffer species are assumed to be at chemical equilibrium. Within each compartment, there are four additional pH equilibrium relations, corresponding to the four buffer pairs. The algebraic relations of the model include the pH equilibria of four buffer pairs,

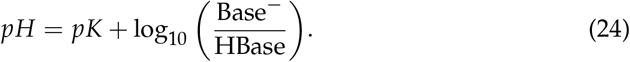

The collection of chosen buffer pairs and even the definition of mass conservation equation can be different over various studies depending on the specific focus of the study. As an example of the variability in these equations, one can compare the presentation of the mass conservation equation for NH_3_:NH_4_^+^ buffer pairs within a cell. In Weinstein’s reports from 1992 and 2009, see [3,14], the authors introduced equation (22) to define the mass conservation equation for these buffer pairs within a cell, while in [4] they used the following equation

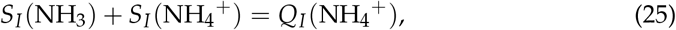

where *Q*_*I*_(NH_4_^+^) is defined as an ammoniagenesis factor, see [4]. According to [3,14], conservation of charge among the buffer reactions requires that

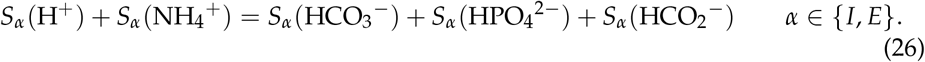

We should mention that in [4], the authors employed a modified version of expression (26) as the requirement of the charge conservation among the cellular buffer reaction in the following form

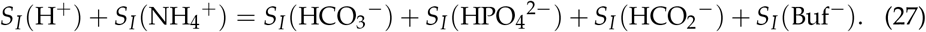

In turn, the charge conservation for the lateral interspace (E) buffer reaction stays unchanged, as given by (26).

Although peritubular PCO_2_ is specified, the CO_2_ concentrations in cell and interspace are model variables. The mass conservation for HCO_3_ ^−^, H_2_CO_3_, CO_2_ is expressed as below

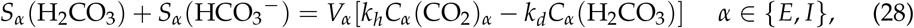

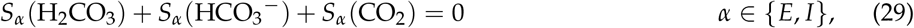

where *k*_*h*_, *k*_*d*_ are the hydration and dehydration rates for CO_2_, respectively. The current study is based on [4], where equation (23) is ignored, and expression (22) is replaced by (25). Also, for the charge conservation among the cellular buffer reaction (26) is replaced by (27). The conservation of mass for the impermeant cellular buffer given by (23) is present in the epithelial model in most of Weinstein’s body of literature. However, we could not see it in [4]. Defining a comprehensive epithelial model which includes all buffer pairs is not straightforward due to the high level of inconsistencies encountered in the literature.

Equations (6),(7),(8),(9),(14) and (17) define a coupled system of 34 differential equations ensuring that mass is conserved^1^.

#### iii. Electroneutrality Constraints

When considering the movement of charged solutes, with valence *Z*_*i*_, *i*∈𝒮, the proposed system of equations is not sufficient to guarantee that the cell and interspace remain electrically neutral. An electroneutrality relation for the cell compartment is expressed through the following balance equation

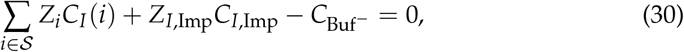

where *C*_Imp_ and 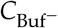 denote the concentration of cell impermeant solute and cell unprotonated buffer, *Z*_*I*,Imp_ is the cell impermeant valence, and *Z*_*i*_ is the valence of species *i, i* ∈𝒮 An electroneutrality relations for the interspace are defined as follows

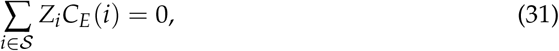

and for all of the buffer reactions, there is conservation of protons, which implies that the following is verified

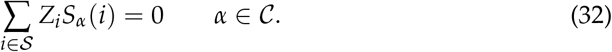

The electroneutrality condition is effectively prescribed by considering that the net charge fluxes into and out of the cell are the same. The membrane charge fluxes can be represented as electrical currents using the following relationships

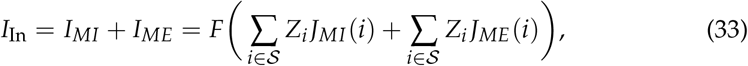

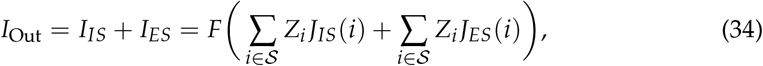

where *F* is Faraday’s constant. Balancing the flow of charge into and out of the cell therefore results in

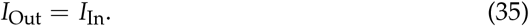

which must hold true at all times. In the current work, according to [4], expression (33) is only considered for the balance of charge transfer across the membranes such that *I*_In_ = 0. Equations (26)-(34) are a collection of different electroneutrality constraints that were introduced in different studies (e.g., [3,4,14]). However, it is important to note that not all these equations were utilised in all different studies. Instead, they were selectively chosen based on the context of each study.

### II. Model Specialisation

The basic principles of mass conservation, pH equilibrium of buffer species, and maintenance of electroneutrality described above apply to epithelial transport in general. To instantiate the general model into a mathematical model for a specific epithelium, all that remains is to define the actual membrane solute and water fluxes of interest to create the specialised model.

#### i. Water Fluxes

With respect to water fluxes, the volume conservation equations for lateral interspace and cell are considered to compute the lateral interspace hydrostatic pressure, and cell volume. Across each cell membrane, the transmembrane water fluxes are proportional to the hydrostatic, oncotic, and osmotic driving forces

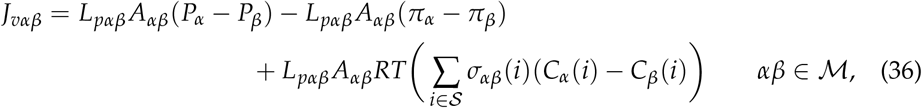

where *P*_*α*_ and *π*_*α*_ are the hydrostatic and oncotic pressures within compartment *α* ∈𝒞, *L*_*pαβ*_ is the membrane water permeability and *σ*_*αβ*_(*i*) is the reflection coefficient of membrane *αβ* ∈ℳ to solute *i* ∈𝒮, and *R* and *T* are the gas constant and absolute temperature, respectively. It is important to mention that all *L*_*pαβ*_ constants which are represented in the implementation of this model (see Python code) are multiplied by *R* and *T*. The reflection coefficients, *σ*_*αβ*_(*i*), stay identical in most of Weinstein’s body of work, see Table 2. In turn, there are some variations in the model parameters (such as coupled transporter coefficients, cell membrane water permeability, and cell membrane solute permeability) across the different works.

#### ii. Convective Solute Fluxes

In the model proposed in [3], it is assumed that there are convective fluxes for all intraepithelial solutes. Considering the definition of the water fluxes, see (36), the convective fluxes are defined as follows [4]

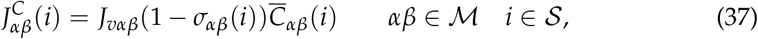

where 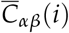 is the logarithmic mean membrane solute concentration described by the expression

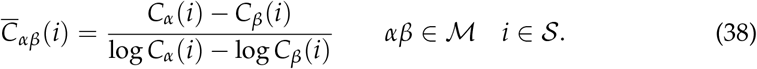

Studying the reflection coefficient values *σ*_*αβ*_(*i*) (defined as membrane-solute properties), one can see that the reflection coefficient is mostly one in lumen-cell (MI), cell-lateral (IE) and cell basal membrane (IS). The most effective membrane to produce the convective fluxes is the interspace basement membrane (ES) with the reflection coefficient mostly zero, and then in the second place it is the tight junction (ME).

#### iii. Passive Solute Fluxes

In the W-PCT-E model, passive solute fluxes across all membranes are assumed to occur by electrodiffusion and to conform to the Goldman-Hodgkin-Katz constant-field flux equation [15]. Passive solute fluxes of the species *i* ∈𝒮across the membrane *αβ* ∈ℳ is therefore given by

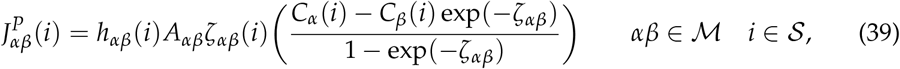

where, for solute *i*∈𝒮 and for membrane *αβ*, ∈ℳ *h*_*αβ*_(*i*) [cm.s^−1^] is the permeability and *ζ*_*αβ*_(*i*) is a normalised electrical potential difference (dimensionless), defined by

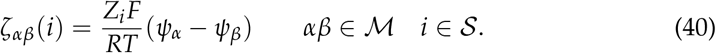

where *ψ*_*α*_ and *ψ*_*β*_ are electrical potentials within compartments *α* and *β*, respectively. *F* is Faraday’s constant, *R* and *T* are the gas constant and absolute temperature, sequentially. Weinstein et al. did not hold on to one solid definition for the permeability, in some cases permeability was multiplied by the area of the corresponding membrane *h*_*αβ*_(*i*)*A*_*αβ*_[10^−5^ cm^3^ .s^−1^ .cm^−2^. epithelium] as an example see Weinstein et al. [4]. For the uncoupled permeation of neutral solutes across membranes, the Fick law is utilised

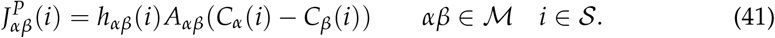

#### iv. Coupled Solute Fluxes

Coupled solute fluxes in the W-PCT-E model include three different categories of transporters: simple cotransporters, simple exchangers, and complex exchangers. All coupled solute transporters in this model have been represented according to linear nonequilibrium thermodynamics, so that the solute permeation rates are proportional to the electrochemical driving force of the aggregate species, with a single permeation coefficient. Simple cotransporters consist of peritubular K^+^ – Cl^−^, luminal Na^+^ – Gluc and Na^+^ – H_2_PO_4_ ^−^, which are in the form of the following equations

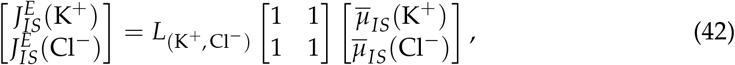

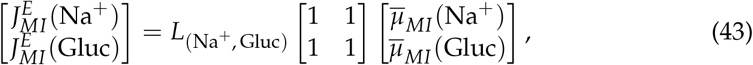

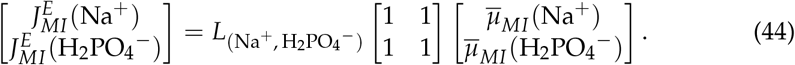

It is important to mention that all transporters within cell-basal (IS) membrane are also considered for the cell-lateral (IE) membrane.

In the equations above, the fluxes of two different species across the cotransporter are equal (1 : 1 stoichiometry).

Simple exchangers such as Na^+^/H^+^, Na^+^/NH_4_^+^, Cl^−^ /HCO_2_ ^−^, and Cl^−^ /HCO_3_ ^−^are located at the lumen-cell (MI) membrane, and represented by the equations below

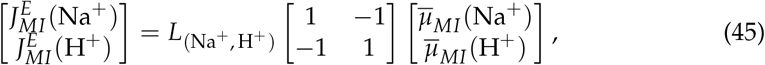

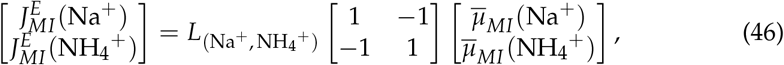

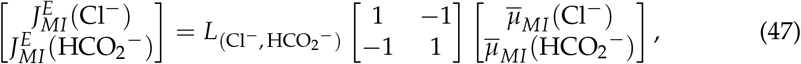

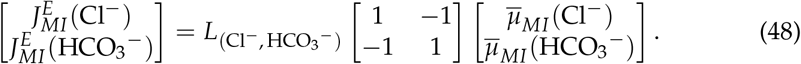

In [3], the authors introduced the NHE3 exchanger in the luminal membrane through equations (45) and (46). However, the NHE3 exchanger has been developed through the kinetic model proposed by Weinstein et al. in 1995 [13]. In the current work, the NHE3 exchanger is modelled by employing the mathematical system introduced in [13]. There are also two more complex transporters at the peritubular membrane: Na^+^ – HCO_3_ ^−^ and Na^+^ – 2 HCO_3_ ^−^ /Cl^−^ which are defined by the following equations (see [3])

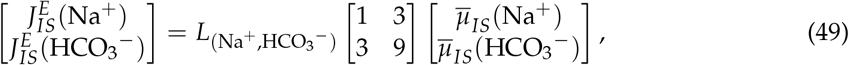

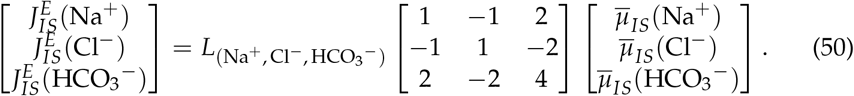

In the expressions above, *L*_(*i,j*)_ is the transporter coupling coefficient, a single proportionality constant which specifies the relative activity of the transporters. In turn, 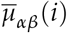 is the electrochemical potential difference of species *i* across all membranes

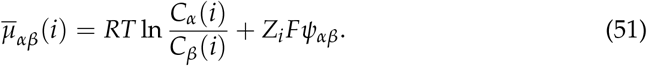

Here, we also define the following quantity

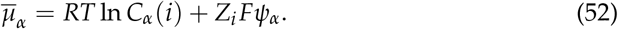

It is important to mention that all permeability coefficients, *L*_(*i,j*)_, which are represented in Table 1, from [3], are scaled (multiplied) by their respective membrane area to take into account the effective behaviour of the representative membrane.

**Table 1.**
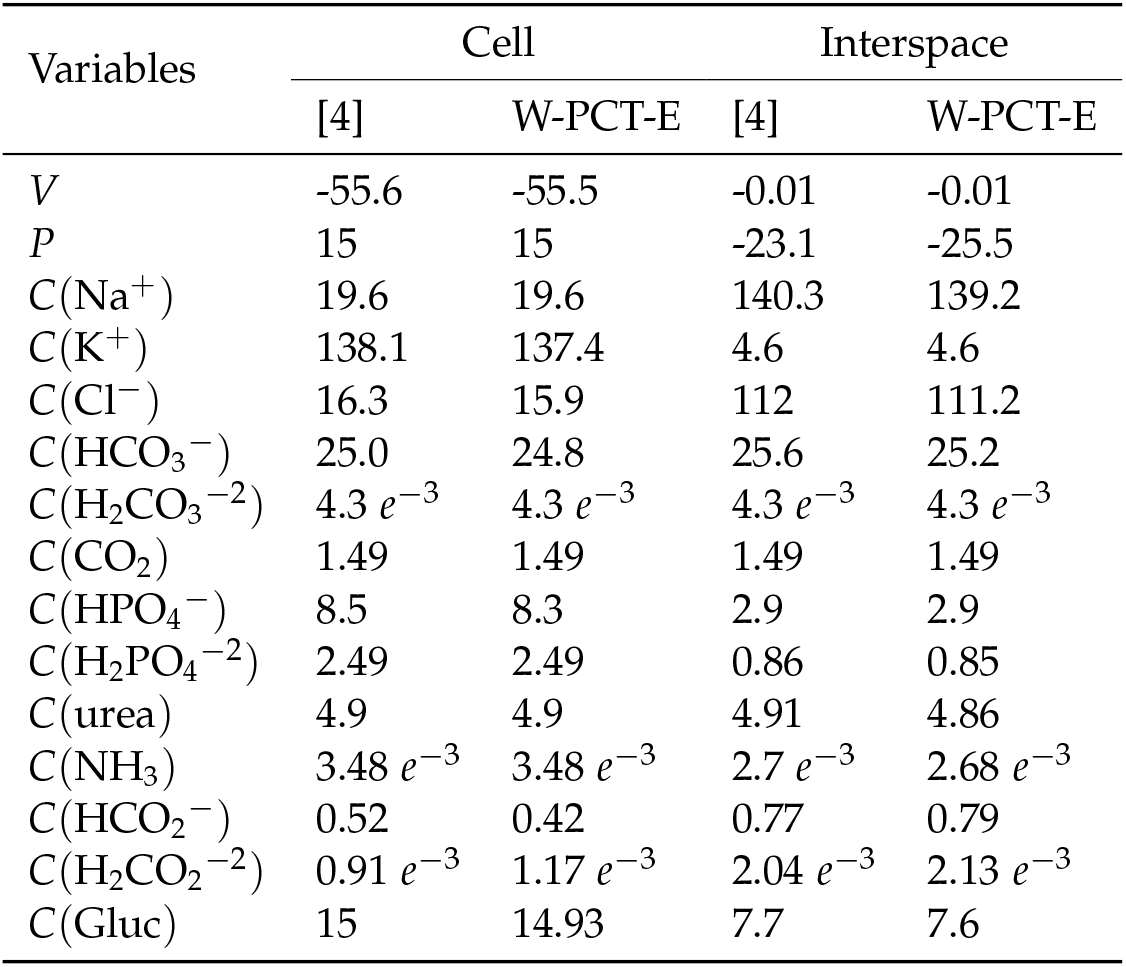
A comparison between the solutions obtained with the present W-PCT-E model and the model reported in [4]. Electrical potentials are in mV, pressures in mmHg and concentrations in M.

| Variables | Cell |  | Interspace |  |
| --- | --- | --- | --- | --- |
|  | [4] | W-PCT-E | [4] | W-PCT-E |
| $V$ | -55.6 | -55.5 | -0.01 | -0.01 |
| $P$ | 15 | 15 | -23.1 | -25.5 |
| $C(\text{Na}^+)$ | 19.6 | 19.6 | 140.3 | 139.2 |
| $C(\text{K}^+)$ | 138.1 | 137.4 | 4.6 | 4.6 |
| $C(\text{Cl}^-)$ | 16.3 | 15.9 | 112 | 111.2 |
| $C(\text{HCO}_3^-)$ | 25.0 | 24.8 | 25.6 | 25.2 |
| $C(\text{H}_2\text{CO}_3^{-2})$ | $4.3 \cdot 10^{-3}$ | $4.3 \cdot 10^{-3}$ | $4.3 \cdot 10^{-3}$ | $4.3 \cdot 10^{-3}$ |
| $C(\text{CO}_2)$ | 1.49 | 1.49 | 1.49 | 1.49 |
| $C(\text{HPO}_4^-)$ | 8.5 | 8.3 | 2.9 | 2.9 |
| $C(\text{H}_2\text{PO}_4^{-2})$ | 2.49 | 2.49 | 0.86 | 0.85 |
| $C(\text{urea})$ | 4.9 | 4.9 | 4.91 | 4.86 |
| $C(\text{NH}_3)$ | $3.48 \cdot 10^{-3}$ | $3.48 \cdot 10^{-3}$ | $2.7 \cdot 10^{-3}$ | $2.68 \cdot 10^{-3}$ |
| $C(\text{HCO}_2^-)$ | 0.52 | 0.42 | 0.77 | 0.79 |
| $C(\text{H}_2\text{CO}_2^{-2})$ | $0.91 \cdot 10^{-3}$ | $1.17 \cdot 10^{-3}$ | $2.04 \cdot 10^{-3}$ | $2.13 \cdot 10^{-3}$ |
| $C(\text{Gluc})$ | 15 | 14.93 | 7.7 | 7.6 |

**Table 2.**

| Parameters | Membrane Properties |  |  |  |  |
| --- | --- | --- | --- | --- | --- |
|  | ME | MI | IE | ES | IS |
| Membrane Area [$\text{cm}^2/\text{cm}^2.\text{epithelial}$] | | | | | |
| $A_{\alpha\beta}$ | 0.001 | 36.0 | 36.0 | Eq:61 | 1.0 |
| Water Permeability $\times RT$ [$\text{cm.s}^{-1}.\text{Osmol}^{-1}$] | | | | | |
| $L_{p\alpha\beta}$ | $0.2e^{+02}$ | $0.2e^{-03}$ | $0.2e^{-03}$ | $0.6e^{+01}$ | $0.2e^{-03}$ |
| Permeability Coefficient, $H_{\alpha\beta}$ [$\text{cm.s}^{-1}$], Eq: 39 | | | | | |
| $\text{Na}^+$ | $0.13 e^{+1}$ | 0.0 | $0.39 e^{-8}$ | $0.5 e^{-1}$ | 1.0 |
| $\text{K}^+$ | $0.145 e^{+1}$ | $0.25 e^{-6}$ | $0.2 e^{-5}$ | $0.7 e^{-1}$ | $0.2 e^{-5}$ |
| $\text{Cl}^-$ | $0.1 e^{+1}$ | 0.0 | 0.0 | $0.6 e^{-1}$ | 0.0 |
| $\text{HCO}_3^-$ | $0.4 e^{+0}$ | $0.1 e^{-7}$ | 0.0 | $0.5 e^{-1}$ | 0.0 |
| $\text{H}_2\text{CO}_3^+$ | $0.4 e^{+0}$ | $0.65 e^{-3}$ | $0.65 e^{-3}$ | $0.5 e^{-1}$ | $0.65 e^{-3}$ |
| $\text{CO}_2$ | $0.4 e^{+0}$ | $0.75 e^{-1}$ | $0.75 e^{-1}$ | $0.5 e^{-1}$ | $0.75 e^{-1}$ |
| $\text{HPO}_4^{-2}$ | $0.2 e^{+0}$ | $0.95 e^{-8}$ | $0.225 e^{-7}$ | $0.4 e^{-1}$ | $0.225 e^{-7}$ |
| $\text{H}_2\text{PO}_4^-$ | $0.2 e^{+0}$ | 0.0 | $0.33 e^{-6}$ | $0.4 e^{-1}$ | $0.33 e^{-6}$ |
| Urea | $0.4 e^{+0}$ | $0.105 e^{-5}$ | $0.1 e^{-5}$ | $0.8 e^{-1}$ | $0.1 e^{-5}$ |
| $\text{NH}_3$ | $0.25 e^{+0}$ | $0.85 e^{-3}$ | $0.1 e^{-2}$ | $0.2 e^0$ | $0.1 e^{-2}$ |
| $\text{NH}_4^+$ | $0.25 e^{+0}$ | $0.215 e^{-6}$ | $0.6 e^{-6}$ | $0.2 e^0$ | $0.6 e^{-6}$ |
| $\text{H}^+$ | $0.3 e^{+2}$ | $0.85 e^{-2}$ | $0.85 e^{-2}$ | $0.3 e^{+2}$ | $0.85 e^{-2}$ |
| $\text{HCO}_2^-$ | $0.7 e^{+0}$ | 0.0 | $0.19 e^{-6}$ | $0.5 e^{-1}$ | $0.19 e^{-6}$ |
| $\text{H}_2\text{CO}_2$ | $0.14 e^{+1}$ | $0.5 e^{-1}$ | $0.6 e^{-1}$ | $0.9 e^{-1}$ | $0.6 e^{-1}$ |
| Glucose | $0.8 e^{-1}$ | 0.0 | $0.75 e^{-5}$ | $0.3 e^{-1}$ | $0.75 e^{-5}$ |
| Reflection Coefficient, $\sigma_{\alpha\beta}$ | | | | | |
| $\text{Na}^+$ | 0.75 | 1.0 | 1.0 | 0.0 | 1.0 |
| $\text{K}^+$ | 0.6 | 1.0 | 1.0 | 0.0 | 1.0 |
| $\text{Cl}^-$ | 0.3 | 1.0 | 1.0 | 0.0 | 1.0 |
| $\text{HCO}_3^-$ | 0.9 | 1.0 | 1.0 | 0.0 | 1.0 |
| $\text{H}_2\text{CO}_3^+$ | 0.9 | 1.0 | 1.0 | 0.0 | 1.0 |
| $\text{CO}_2$ | 0.9 | 1.0 | 1.0 | 0.0 | 1.0 |
| $\text{HPO}_4^{-2}$ | 0.9 | 1.0 | 1.0 | 0.0 | 1.0 |
| $\text{H}_2\text{PO}_4^-$ | 0.9 | 1.0 | 1.0 | 0.0 | 1.0 |
| Urea | 0.7 | 0.95 | 0.95 | 0.0 | 0.95 |
| $\text{NH}_3$ | 0.3 | 0.5 | 0.5 | 0.0 | 0.5 |
| $\text{NH}_4^+$ | 0.6 | 1.0 | 1.0 | 0.0 | 1.0 |
| H | 0.2 | 1.0 | 1.0 | 0.0 | 1.0 |
| $\text{HCO}_2^-$ | 0.3 | 1.0 | 1.0 | 0.0 | 1.0 |
| $\text{H}_2\text{CO}_2$ | 0.7 | 0.95 | 0.95 | 0.0 | 0.95 |
| Glucose | 1.0 | 1.0 | 1.0 | 0.0 | 1.0 |
| Coupled Transport Pathways: [$\text{mmol}^2.\text{J}^{-1}.\text{s}^{-1}.\text{cm}^{-2}$] | | | | | |
| $\text{K}^+.\text{Cl}^-$ | - | - | $0.5 e^{-8}$ | - | $0.5 e^{-8}$ |
| $\text{Na}^+.\text{2 HCO}_3^-/\text{Cl}^-$ | - | - | $0.14 e^{-6}$ | - | $0.14 e^{-6}$ |
| $\text{Na}^+.\text{3 HCO}_3^-$ | - | - | $0.15 e^{-7}$ | - | $0.15 e^{-7}$ |
| $\text{Na}^+/\text{H}^+$ | - | $0.225 e^{-7}$ | - | - | - |
| $\text{Na}^+/\text{NH}_4^+$ | - | $0.15 e^{-8}$ | - | - | - |

Table 2 – continued from previous page
| Parameters | Membrane Properties |  |  |  |  |
| --- | --- | --- | --- | --- | --- |
|  | ME | MI | IE | ES | IS |
| Na<sup>+</sup>-glucose | - | $0.75 e^{-8}$ | - | - | - |
| Na<sup>+</sup>-H<sub>2</sub>PO<sub>4</sub><sup>-</sup> | - | $0.5 e^{-8}$ | - | - | - |
| Cl<sup>-</sup>/HCO<sub>2</sub><sup>-</sup> | - | $0.5 e^{-8}$ | - | - | - |
| Cl<sup>-</sup>/HCO<sub>3</sub><sup>-</sup> | - | $0.2 e^{-8}$ | - | - | - |
| Active Transport: [mmol<sup>2</sup>.J<sup>-1</sup>.s<sup>-1</sup>.cm<sup>-2</sup>] |  |  |  |  |  |
| H<sup>+</sup> | - | Eq: 53 | - | - | - |
| Na<sup>+</sup>K<sup>+</sup>-ATPS | - | - | Eq:58 | - | Eq:58 |
| Parameters | Compartment Properties |  |  |  |  |
|  | M | E | I | S |  |
| Concentration, $C_{\alpha}(i)$[mol/l] & Valence Number, $Z_i$ | | | | | |
| Na<sup>+</sup> | 0.14 | 0.14 | 0.02 | 0.14 | 1 |
| K<sup>+</sup> | $0.49 e^{-2}$ | $0.46 e^{-2}$ | 0.13 | $0.49 e^{-2}$ | 1 |
| Cl<sup>-</sup> | 0.113 | 0.11 | $0.16 e^{-1}$ | 0.113 | -1 |
| HCO<sub>3</sub><sup>-</sup> | 0.024 | 0.25 | $0.25 e^{-1}$ | 0.024 | -1 |
| H<sub>2</sub>CO<sub>3</sub> | $0.44 e^{-5}$ | $0.4 e^{-5}$ | $0.43 e^{-5}$ | $0.44 e^{-5}$ | 0 |
| CO<sub>2</sub> | $0.15 e^{-2}$ | $0.14 e^{-2}$ | $0.14 e^{-2}$ | $0.15 e^{-2}$ | 0 |
| HPO<sub>4</sub><sup>-2</sup> | $0.2 e^{-2}$ | $0.86 e^{-3}$ | $0.94 e^{-2}$ | $0.2 e^{-2}$ | -2 |
| H<sub>2</sub>PO<sub>4</sub><sup>-</sup> | $0.927 e^{-3}$ | $0.28 e^{-2}$ | $0.27 e^{-2}$ | $0.927 e^{-3}$ | -1 |
| CH<sub>4</sub>N<sub>2</sub>O | $0.5 e^{-2}$ | $0.4 e^{-2}$ | $0.4 e^{-2}$ | $0.5 e^{-2}$ | 0 |
| NH<sub>3</sub> | $0.282 e^{-5}$ | $0.26 e^{-5}$ | $0.35 e^{-5}$ | $0.282 e^{-5}$ | 0 |
| NH<sub>4</sub><sup>+</sup> | $0.197 e^{-3}$ | $0.17 e^{-3}$ | $0.23 e^{-3}$ | $0.197 e^{-3}$ | 1 |
| HCO<sub>2</sub><sup>-</sup> | $0.1 e^{-2}$ | $0.7 e^{-3}$ | $0.5 e^{-3}$ | $0.1 e^{-2}$ | -1 |
| H<sub>2</sub>CO<sub>2</sub> | $0.285 e^{-6}$ | $0.2 e^{-6}$ | $0.9 e^{-7}$ | $0.285 e^{-6}$ | 0 |
| C<sub>6</sub>H<sub>12</sub>O<sub>6</sub> | $0.5 e^{-2}$ | $0.7 e^{-2}$ | $0.1 e^{-1}$ | $0.5 e^{-2}$ | 0 |
| Impermeant | 0.0 | 0.0 | Variable, Eq. (60) | 0.002 | -1 |
| Osmolality | 298.8 | 304 | 301 | 298.8 |  |
| Electrical Potential [mV] |  |  |  |  |  |
| $\psi_{\alpha}$ | Variable, Eq. (39) | Variable | Variable | 0.0 | |
| Hydrostatic Pressure [mmHg] |  |  |  |  |  |
| $P_{\alpha}$ | 15.0 | Variable, Eq. (37) | 15 | 0.0 | |
| Volume [cm<sup>3</sup>/cm<sup>2</sup>. epithelial] |  |  |  |  |  |
| $V_{\alpha}$ | Variable Eq. (62) | Variable Eq. (60) | - | - | |

The table can be categorised to two different parts: first, membrane properties (ME, MI, IE, ES,IS); then, compartment properties (M, E, I, S).

**Table 3.** W-PCT-E Model’s Constant Parameters.

| Constant Parameters in PCT-Epithelial model |  |  |  |
| --- | --- | --- | --- |
| Constant | Definition | Unit | Value |
| $R$ | Gas Const | J/mol.K | 8.314 |
| $T$ | Temp | K | 273.15 |
| $F$ | Faraday | C/mol | $96.5 e^3$ |
| $K_h$ | Hydration Const $\text{CO}_2$ | 1/s | $1.45 e^3$ |
| $K_d$ | Dehydration Const $\text{CO}_2$ | 1/s | $4.96 e^5$ |
| $\eta$ | Fluid Viscosity | mmHg/s | $6.4 e^{-6}$ |
| $P(\text{HCO}_3^-)$ | $pK$ of $\text{HCO}_3^-$ | — | 3.57 |
| $P(\text{HCO}_2^-)$ | $pK$ of $\text{HCO}_2^-$ | — | 3.76 |
| $P(\text{NH}_3)$ | $pK$ of $\text{NH}_3$ | — | 9.15 |
| $P(\text{HPO}_4^-)$ | $pK$ of $\text{HPO}_4^{-2}$ | — | 6.8 |
| $P(\text{Buf}^-)$ | $pK$ of Cell Buffers | — | 7.5 |
| $V_{I0}$ | Reference Cell Volume | $\text{cm}^3/\text{cm}^2.\text{epithelium}$ | $0.1 e^{-2}$ |
| $C_{\text{IMP0}}$ | Reference Cell Impermeant | mol/l | $0.6 e^{-1}$ |
| $C_{\text{TBUF}}$ | Total Cell Buffer Conc | mmol/ml | $0.6 e^{-1}$ |
| $Q_{\text{NH}_4^+}$ | Ammonia generation | mmol/s.$\text{cm}^2$ | $0.7e - 7$ |

Here, $pK = \log(\frac{1}{K_\alpha})$ and $K_\alpha$ represents the equilibrium constant for different buffer pairs. 654
In some cases, we use $RT = 1.93 e^4 \text{ mmHg.ml/mmol}$ and in some other cases $RT = 2.57 \text{ J/mmol}$ just to simplify the calculations. 655 656 657
We need to highlight that in here $C_{\text{TBUF}} = C_{\text{BUF}} + C_{\text{HBUF}}$ , $C_{\text{BUF}}$ and $C_{\text{HBUF}}$ are variables and they indicate the cell buffer concentration and protonated cell buffer concentration, respectively. 658 659 660

#### v. Active Solute Fluxes

In the W-PCT-E model, there are two ATPases, the apical membrane H^+^ – ATPase and a peritubular Na^+^/K^+^ – ATPase. To model the H^+^ – ATPase, an expression of the following form is utilised

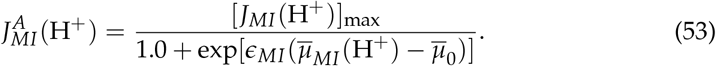

Note that the rate of proton pumping varies as a function of the transmembrane electrochemical potential difference. Here, [*J*_*MI*_ (H^+^)]_max_ is a maximal rate of transport, and *e*_*MI*_ is a steepness coefficient. The Na^+^/K^+^ – ATPase exchanges three cytosolic Na^+^ ions for two peritubular cations, K^+^ or NH_4_^+^, in the way that compete for the binding. The following expressions represent all the three different fluxes due to the Na^+^/K^+^ – ATPase activities

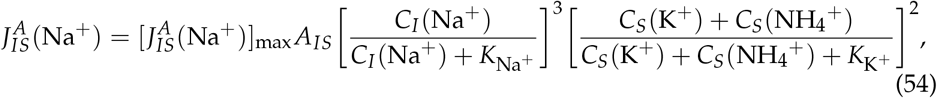

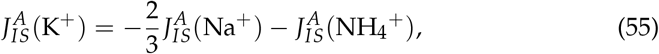

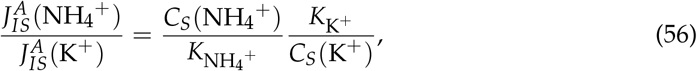

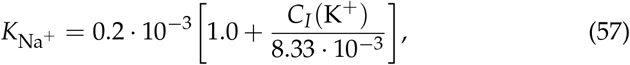

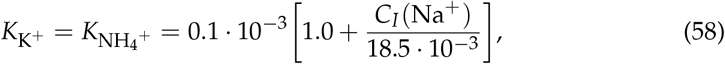

in which *K*_Na_^+^, the half-maximal of Na^+^ concentration, which scales linearly with the cellular concentration of K^+^; *K*_K_^+^, which is the half-maximal of K^+^ concentration, rises linearly with the external concentration of K^+^. The pump flux of K^+^ plus NH_4_^+^ reflects the 3:2 stoichiometry. Similar expressions are considered for active transport at the cell-lateral membrane (IE), denoted by 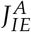. Figure 3 illustrates the proximal PCT cell featuring coupled transport pathways and ion channels within luminal and basolateral membranes, the coupled transport pathways within the cell-lateral are not included.

**Figure 3.**
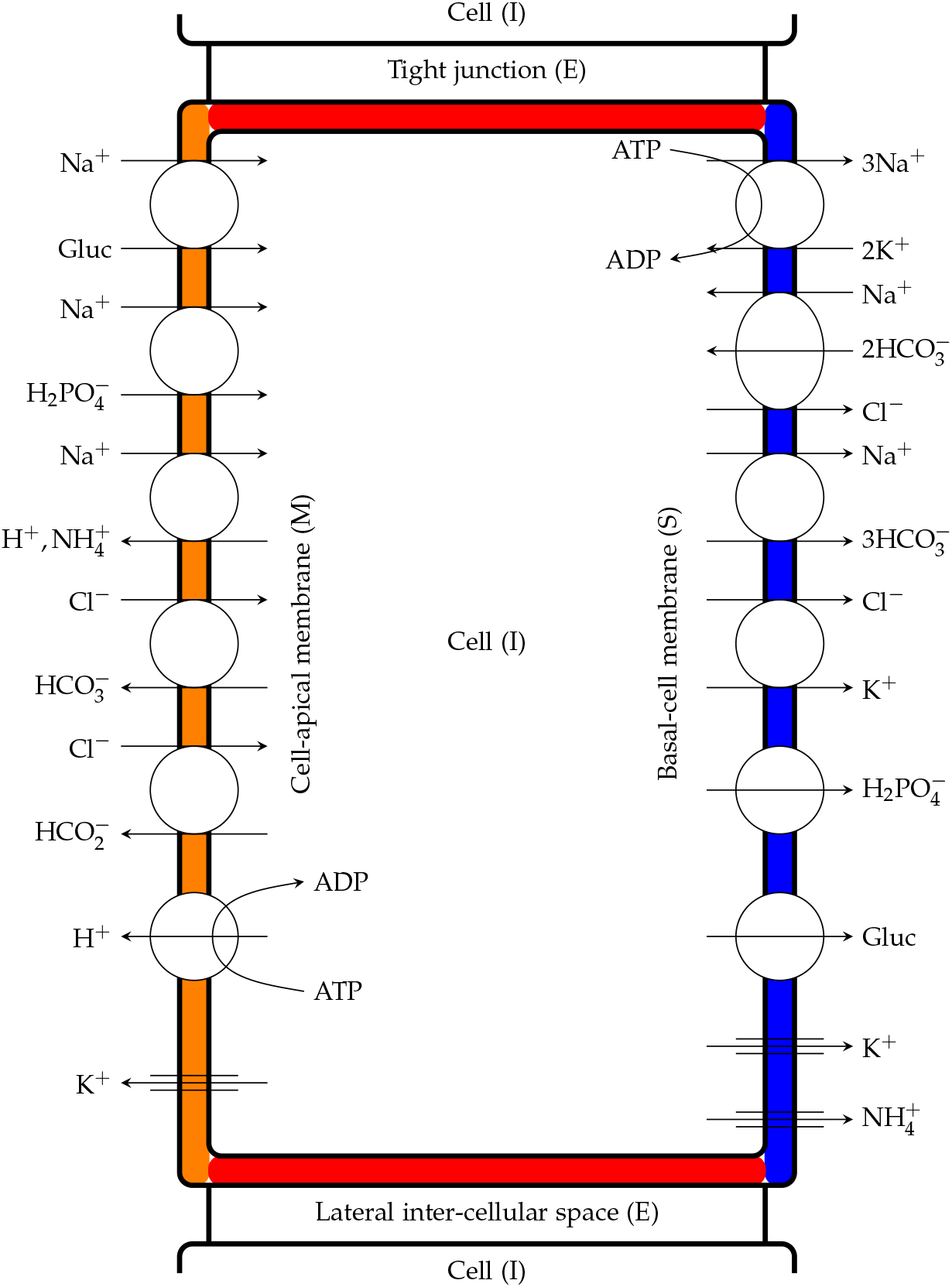
Proximal tubule cells showing coupled transport pathways and some ion channels within the luminal and peritubular cell membranes. It is essential to mention that peritubular cell and cell-lateral membranes share the same feature regarding their transport pathways, even though it has not been shown in this diagram.

#### vi. Total Membrane Solute Fluxes

In summary, in the W-PCT-E model, the intraepithelial solute transport through membrane *αβ* ∈ M results from the contribution of convective flux 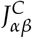, passive flux 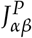, electrodiffusive flux 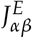, and/or metabolically driven flux 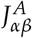, that is

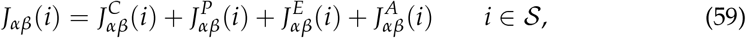

where *J*_*αβ*_(*i*)[mmol.s.cm^−2^] is the total flux of *i* solute flow from the compartment *α* to the compartment *β* (through membrane *αβ*).

#### vii. Model compliance parameters

In the W-PCT-E model, the cell is compliant in a manner that there is no hydrostatic pressure difference between the cell and lumen, therefore, we have *P*_*I*_ = *P*_*M*_. There is a substantial oncotic force within the cell, *π*_*I*_, that increases with decrements in the cell volume. Here, it is assumed that the total cell protein content *C*_*I*,Imp_*V*_0*I*_ is fixed and that *π*_*I*_ is proportional to *C*_*I*,Imp_, for this reason *C*_*I*,Imp_ replaces *π*_*I*_ as one of the model unknowns (for more information see [3,4])

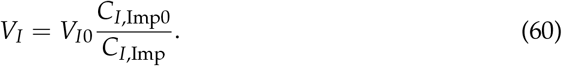

The lateral interspace volume (*V*_*E*_) and its basement membrane area (*A*_*ES*_) are functions of interspace hydrostatic pressure, *P*_*E*_, that is

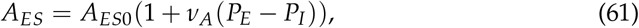

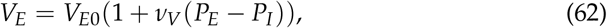

where *A*_*ES*0_ and *V*_*E*0_ are reference values for outlet area and volume, *ν*_*A*_ and *ν*_*V*_ are the compliance parameters used to ensure that suitable pressures are maintained in the system (see Table 2). To define the epithelial thickness or epithelial volume [cm^3^/cm^2^. epithelium], the cell volume and channel volume are added as below,

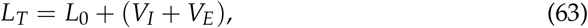

where *L*_*T*_ is also considered as an epithelial height. As one can see, the channel (LIS) volume is a function of intracellular pressure. Table 2 includes the model parameters and their definitions either in the current document or in the Python code. You can see the geometric parameters for all compartments in the tables reported below. As one can see, the luminal and peritubular cell membranes have equal areas, i.e., 36 [cm^2^/cm^2^. epithelium]. The lateral interspace is compliant and distends with transport-associated increments in interspace pressure. In this model, the tight junction properties are fixed and do not vary with transjunctional pressure differences.

#### III. Model Calculations

In this work, the W-PCT-E model is bathed on both luminal and peritubular sides by solutions of equal concentration. Baseline bath and lumen conditions are those reported in Table 2. Choices for model parameters appear in Table 2. The geometric parameters are completely different from [3], *A*_*IE*_= 35 [cm^2^/cm^2^. epithelium], *A*_*IM*_ = 36 [cm^2^/cm^2^. epithelium]. In the LIS interspace, The LIS basement membrane area and volume are compliant and distend with transport-associated increments in interspace pressure. However, the membrane areas in the cell are fixed and do not vary with transjunctional pressure differences. The cell volume varies linearly as a function of the cellular impermeant concentration. The suitability of these parameters was not tested here. For the W-PCT-E simulations, the 35 nonlinear ordinary differential equations are solved using a finite difference numerical method for time discretisation along the Python solver “scipy.optimize.root”. The evaluation of the model involves integrating the mass conservation equations from an initial time to a final time using small time increments. Simulation time is chosen so that we ensure that steady-state regime is reached. Here, there is a list of all variables in the model: First, all variables appear in the lateral interspace compartment, *V*_*E*_, *P*_*E*_, *C*_*E*_(Na^+^), *C*_*E*_(K^+^), *C*_*E*_(Cl^−^), *C*_*E*_(HCO_3_ ^−^), *C*_*E*_(H_2_CO_3_ ^− 2^), *C*_*E*_(CO_2_), *C*_*E*_(HPO_4_ ^−^), *C*_*E*_(H_2_PO_4_ ^− 2^), *C*_*E*_(urea), *C*_*E*_(NH_4_), *C*_*E*_(HCO_2_ ^−^), *C*_*E*_(H_2_CO_2_ ^− 2^), *C*_*E*_(Gluc).

Then, all variables which appear in the cellular compartment, *V*_*I*_, *C*_*I,Imp*_, *C*_*I*_(Na^+^), *C*_*I*_(K^+^), *C*_*I*_(Cl^−^), *C*_*I*_(HCO_3_ ^−^), *C*_*I*_(H_2_CO_3_ ^− 2^), *C*_*I*_(CO_2_), *C*_*I*_(HPO_4_ ^−^), *C*_*I*_(H_2_PO_4_ ^− 2^), *C*_*I*_(urea), *C*_*I*_(NH_4_), *C*_*I*_(HCO_2_ ^−^), *C*_*I*_(H_2_CO_2_ ^− 2^), *C*_*I*_(Gluc), *C*_*I*_(Buf^−^), *C*_*I*_(HBuf), plus the only luminal variable which is the voltage inside the lumen *V*_*M*_. The Github link for the W-PCT-E Python code is https://github.com/iNephron/W-PCT-E.

## Results

We investigated the W-PCT-E model validity by designing several experiments; the analyses are performed over the steady-state solutions found from numerical simulations. To test the W-PCT-E model robustness, we investigated the sensitivity of steady-state solutions to different sets of initial conditions or the time-steps. Furthermore, we explored reproducibility by replicating some simulation experiments reported in [3,4] using the W-PCT-E. We then investigated the sensitivity to salt intake or luminal salt concentration in the W-PCT-E model based on earlier work [16]. Structural analyses were performed by inhibiting the key transporters in different membranes, such as the Na^+^/K^+^-ATPase in the peritubular membrane or SGLT, NHE3, and Na^+^-H_2_PO_4_ transporters in the apical membrane, and relating the predicted responses to observed biological phenomena.

### i. Model Sensitivity Analysis

As reported in [17], a time step size no greater than Δ*t* = 0.1 s is required to ensure a converged numerical solution for epithelial transport models.

In exploring the sensitivity of our W-PCT-E model to the initial conditions, we found that as long as the initial conditions were within a reasonable physiological range that the steady-state solution was insensitive to the initial conditions (to check the initial conditions, see the https://github.com/iNephron/W-PCT-E). Outside that range, however, the model results are highly variable. To account for this, we follow a simulation protocol whereby we initially disable the active transporters and allow the system to reach an initial steady state using only the passive water transport. This ensures that the simulations then begin at a suitable initial state to which the predicted steady-state solution is insensitive.

### ii. Model Reproducibility

The present W-PCT-E model is built from a collection of mathematical representations reported in the literature. Our efforts have been to compile model parameters and equations from different scientific studies in which the components of PCT have been reported. Moreover, we provide the community with a freely available implementation to speed up research. In doing so, we aim to ensure that the behaviour of the W-PCT-E model reproduces that of the source model. Here, we provide exemplars exhibiting the flexibility and reproducibility of the W-PCT-E model.

#### Flow-dependent transport in the PC

The mathematical model of the rat proximal tubule [4] was designed to include the calculation of microvillous torque and to incorporate torque-dependent solute transport in a compliant tubule. Here, we aim to reproduce some of the results reported in [4] by tuning the parameters according to [4] (see Tables 1-2 in that article). For those constant parameters or boundary conditions which were not defined in [4], we infer them from earlier works, specifically [2,3]. Reproducing these simulation experiments, we found no significant discrepancies, as shown in Table 1, for steady-state solutions such as solute concentrations, luminal voltage, and luminal pressure.

#### Chloride transport in the PCT

Here, we aim to reproduce some of the results from chloride transport in the proximal tubule [3] to explore the possible interactions between the individual transporter pathways and their contribution to overall chloride reabsorption in a proximal tubule. At the apical membrane, Cl^−^ /HCO_2_ ^−^ and Cl^−^ /HCO_3_ ^−^ exchangers are the main pathways for Cl^−^ entry, and across the peritubular membrane Cl^−^ /2HCO_3_ ^−^ -Na^+^ and Cl^−^ -K^+^ for the exit of Cl^−^. At the tight junction, chloride fluxes are both diffusive and convective. We performed the same experiments as reported in [3] and observed that the W-PCT-E model (with parameters modified accordingly) predicted similar behaviour as [3].

Figure 4(a) represents the effect of luminal HCO_3_ ^−^ concentration on the cellular, tight junction, and total epithelial Cl^−^ fluxes; the total and junctional fluxes illustrate a dramatic decrease in comparison to the cellular flux, which depicts a modest increase. In panel (b), one can see that the general effect of HCO_2_ ^−^ concentration on overall Cl^−^ absorption for all the fluxes is relatively modest. Panel (c) displays the impact of HCO_2_ ^−^ concentration, while the luminal and peritubular cell membrane permeability for H_2_CO_2_ was set at 10% of the original value in panel (a) and (b). To maximise the impact of the luminal and peritubular HCO_2_ ^−^ concentrations for the case of the small H_2_CO_2_ permeability, in [3] the authors assumed that all luminal Cl^−^ entry is through Cl^−^ /HCO_2_ ^−^ exchange. The W-PCT-E demonstrates that small luminal and peritubular cell membrane permeability for H_2_CO_2_ could not sustain any substantial luminal Cl^−^ flux. These results confirm previously published findings according to [3,18].

**Figure 4.**
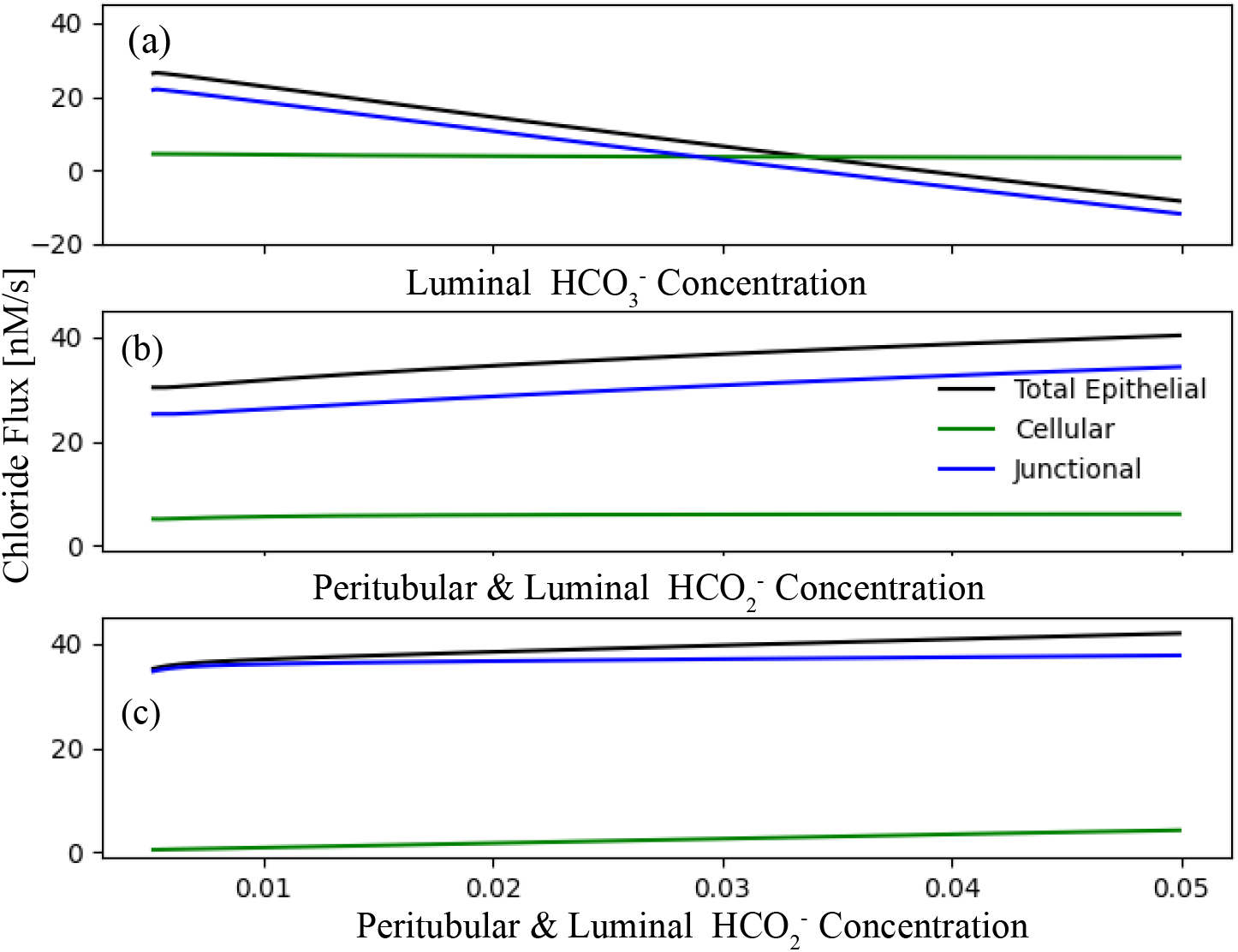
Total epithelial chloride fluxes for the W-PCT-E model. Panel (a) represents the effect of luminal HCO_3_ ^−^ concentration on Cl^−^ fluxes (c.f. Fig. 9, [3]). Panel (b) represents the effect of ambient HCO_2_ ^−^ concentration on Cl^−^ fluxes (c.f. Fig. 11, [3]). Luminal concentration for HCO_3_ ^−^ is considered to be as low as 0.004 M, while the peritubulur concentration for HCO_3_ ^−^ stays the same 0.024 M. Luminal and peritubular formate are varied simultaneously from 0.003 to 0.05. Panel (c) shows the effect of ambient HCO_2_ ^−^ concentration on Cl^−^ fluxes (c.f. Fig. 12, [3]). Here, the luminal Cl^−^ entry proceeds exclusively through Cl^−^ /HCO_2_ ^−^ exchanger, which means the coupled transport coefficient for Cl^−^ /HCO_3_ ^−^ is considered to be zero. Also, apical and peritubular membrane permeabilities are set at 10% of original reference in Table 2. Luminal concentration for HCO_3_ ^−^ is considered to stay at 0.004 M, while HCO_3_ ^−^ peritubular concentration is set to 0.024 M.

#### Salt Sensitivit

Next, we test the flexibility, reusability, and reproducibility of the W-PCT-E model by reproducing a simple model of Na^+^ transport in the mammalian urinary bladder to study the salt sensitivity [16]. Here, we design the same experiment where the Na^+^ and Cl^−^ concentrations are increased in a step-wise manner. At each step, we performed an 8% increase of both luminal and peritubular bath concentrations (see Figure 5(a)). This resulted in the step-wise increase in the cellular activities for the primary solutes, which are displayed in Figure 5(c,d,e). The changes in voltage result from step-wise increase in the bathing solution activities of Na^+^ and Cl^−^ ; Figure 5(b) shows that there is a decrease in voltage after each sequential increase in NaCl. One can see that there is a good agreement between the W-PCT-E model and the Na^+^ transport model in the mammalian urinary bladder reported in [16,19]. We should mention that in the studies cited above, there are only three main primarily solutes (Na^+^, K^+^ and Cl^−^) and one active transporter, the Na^+^/K^+^-ATPase.

**Figure 5.**
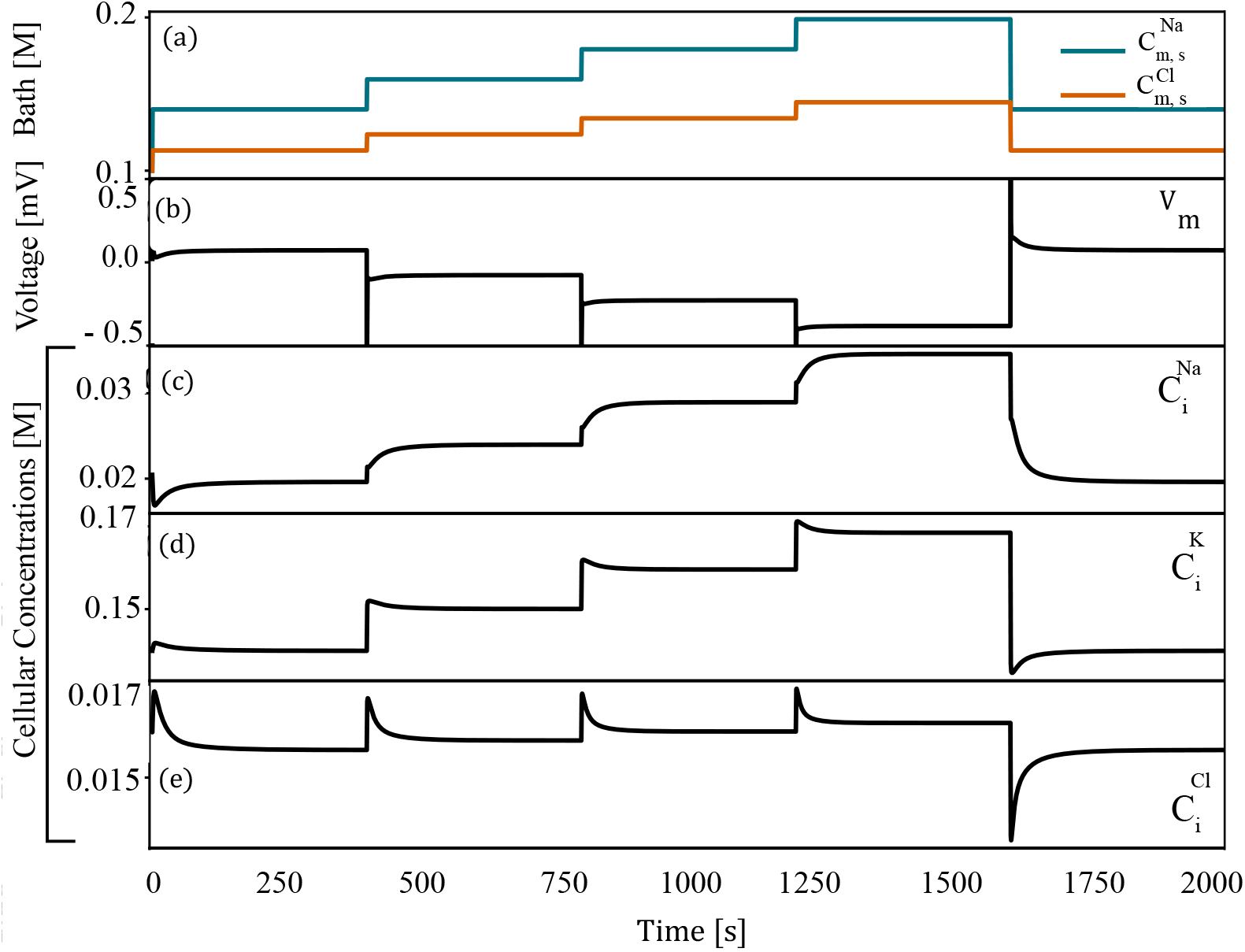
Effect of changes of luminal and peritubular Na^+^ and Cl^−^ concentrations on the cellular system, c.f., Figures 8 and 9 in [16]. Panel (a): both peritubular and luminal sodium and chloride concentrations are increased in a step-wise manner, at each step there is an increase of 8% in regard to the baseline bath conditions. Panel (b): transepithelial potential changes due to the changes in sodium and chloride concentrations. Panels (c)-(e): changes on some selected cellular solutes concentrations.

### iii. Structural Analysis

The goal of the present study was also to create and make available a mathematical model of epithelial transport that is sufficiently flexible to accommodate the investigation of different physiological phenomena in the epithelial system. Here we performed a structural analysis of the W-PCT-E model to both demonstrate this flexibility and to explore the application of this model to a range of physiological perturbations.

To investigate the effect of each transporter in the W-PCT-E model on the overall behaviour, we performed experiments in which we individually inhibited each of the transporters and compared the total epithelial fluxes. We illustrate some of these results in the following section; the first set of simulations addresses the inhibition of both basal and cell-lateral transporters and the second set of simulations addresses the inhibition of the apical cell transporters.

#### iii.1 Inhibition of Peritubular (IS and IE) Transporters

We separately eliminated the Na^+^/K^+^-ATPase and two symporters (K^+^-Cl^−^ and Na^+^-HCO_3_ ^−^) on both the cell-basal and cell-lateral membranes and observed the resulting membrane fluxes and cellular concentrations. Inhibition of each transporter was accomplished by setting the coupling transport coefficient to zero.

We present our results in Figure 6. Panel (a) displays the membrane fluxes (ES, IE, IS, ME, MI) and cellular concentrations for the four primary solutes (Na^+^, K^+^, Cl^−^, Glucose) in the case of the original full W-PCT-E model. Panel (b) represents the results when considering the inhibition of the Na^+^/K^+^-ATPase, from which one can observe a reduction in all the membrane fluxes and notable changes in the cellular solute concentrations; see bottom row in Figure 6(b). There is a considerable reduction in Na^+^ membrane fluxes, which confirms the critical role of the Na^+^/K^+^-ATPase in the production of Na^+^ fluxes. Inhibition of the pump stops sodium exit and potassium entry into the cell; thereby, sodium concentration increases (from 19.6 mmol/L to 142.0 mmol/L) while there is a decrease in the cellular potassium concentration (from 137.3 mmol/L to 8.4 mmol/L).

**Figure 6.**
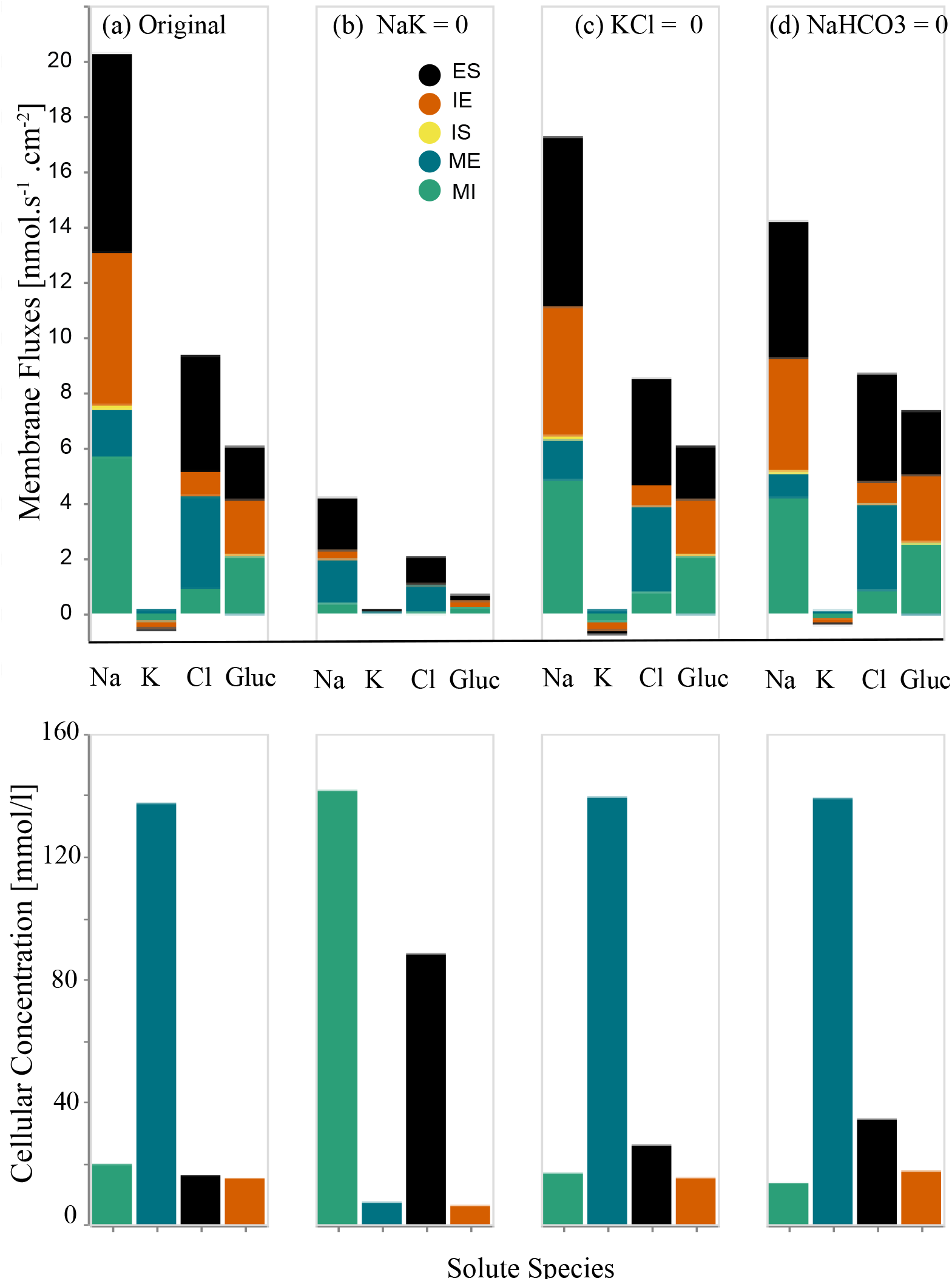
Changes in the membrane fluxes and cellular concentrations due to the inhibition of transporters on the cell-basal and cell-lateral membranes. First row: in each panel, we present four sets of results for four different configurations that depict the total membrane fluxes for the following species: Na^+^, K^+^, Cl^−^, Glucose. The total membrane fluxes include all the membrane activities from five membranes, IS, ME, MI, IE, ES, which are stacked on top of each other. Panel (a) represents the original full model (control configuration). Panel (b) represents scenario due to the Na^+^-K^+^ pump elimination. Panel (c) corresponds to the scenario of K^+^-Cl^−^ elimination, panel (d) is for the inhibition of Na^+^-HCO_3_ ^−^ transporters. Second row: we illustrate the cellular concentrations corresponding to the related configuration for the same species: Na^+^, K^+^, Cl^−^, Glucose.

Figure 6(c) illustrates the effect of the inhibition of K^+^-Cl^−^ transporter on the W-PCT-E model activity. One can observe a decrease in total fluxes both for sodium and chloride. While there is a decrease in the sodium concentration, there is an increase in potassium and chloride concentrations. Figure 6(d) illustrates the response of the W-PCT-E model in the case of elimination of Na^+^-HCO_3_ ^−^ transporter. There is a significant decrease in sodium total flux, accompanied by notable growth in glucose fluxes. While there is a decrease in the sodium concentration, one can see an increase in other solute concentrations. To better understand the underlying mechanisms of the W-PCT-E model responses to these structural changes, we narrow our focus to sodium. We then study how the elimination of Na^+^/K^+^-ATPase can affect the sodium activity on the epithelial membrane. We need to clarify that there are no transporters on the tight junction and interspace basement and the corresponding fluxes in these membranes are either convective or passive. While in the cell-basal, cell-lateral, and apical cell membrane, the passive and convective fluxes are negligible, for the reflection and permeability coefficient values are considered to be very small, see Table 1. The electrochemical fluxes represent the primary source of fluxes in the cell-basal, cell-lateral, and apical cell membranes.

Figure 7(a) features all different sodium membrane fluxes. In Figure 7(b), we narrow our focus down to sodium fluxes on the epithelial membrane (summation of the tight junction and apical cell membrane activities) and its components. The epithelial activities subdivide into three components: convective, passive, and electrochemical activities. In panel 7(c), we subdivide the electrodiffusive activities into their segments, which are NHE3, SGLT, and Na^+^-H_2_PO_4_ ^−^ transporters.

**Figure 7.**
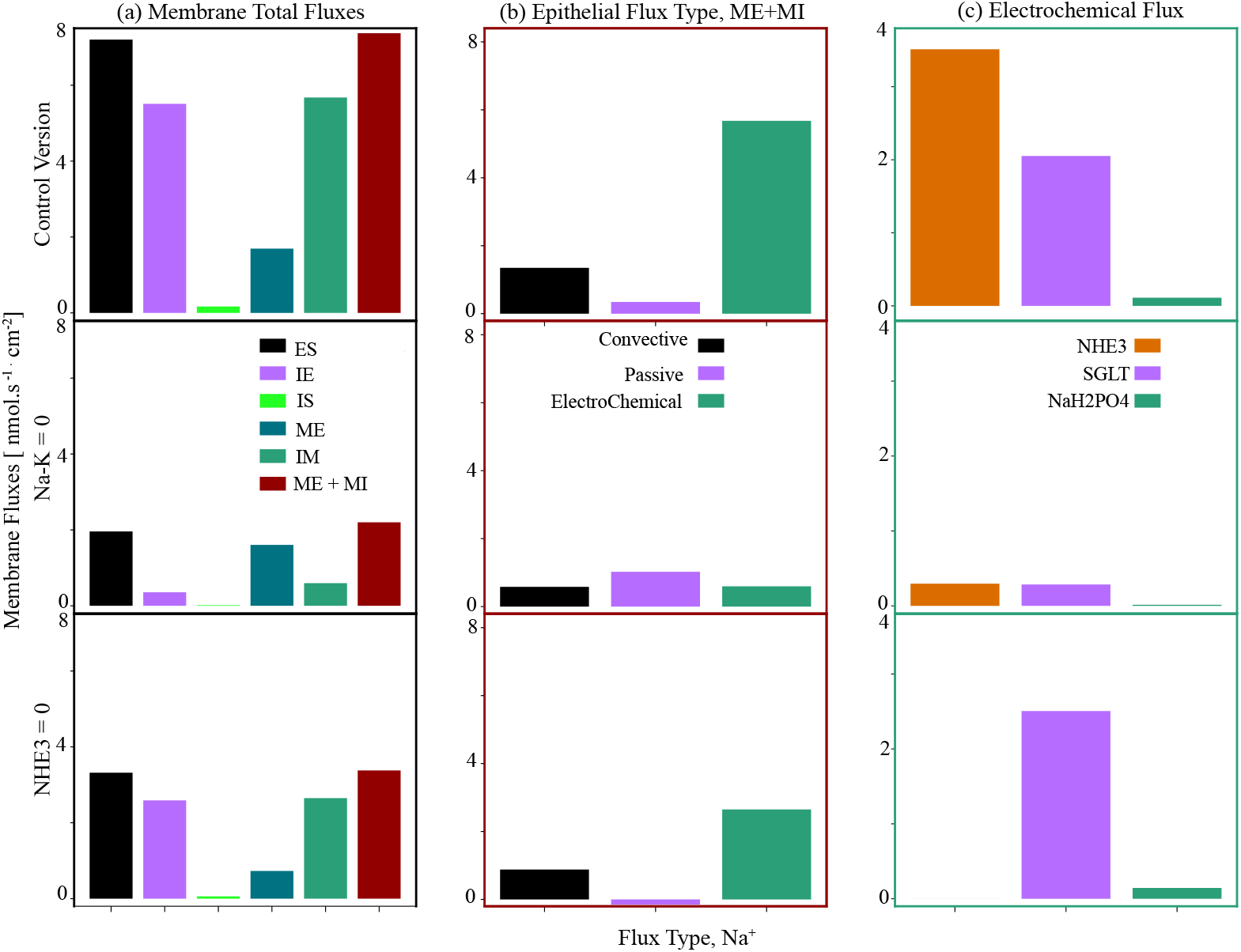
Total epithelial sodium fluxes and the contribution of various sodium flux types. Panel (a) illustrates the different membrane fluxes. Panel (b) presents the epithelial sodium fluxes classified into convective, passive and electrochemical types. Panel (c) details the electrodiffusive activities into their segments: NHE3, SGLT, and 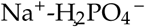. The first row corresponds to the full (control) model, the second row is for the case of elimination of the Na^+^/K^+^-ATPase, and in the third row the results are for the elimination of NHE3.

The first row in Figure 7 represents the sodium fluxes for the full W-PCT-E model, considered as the control version. In the second row, we illustrate the sodium fluxes due to the elimination of the Na^+^/K^+^-ATPase. By a simple comparison between the first and second rows, one can see that the membrane activities drop considerably after the Na^+^/K^+^-ATPase inhibition; as an example, the total epithelial sodium flux decreases from 7.39 nmol.s^-1^.cm^-2^ to 2.19 nmol.s^-1^.cm^-2^, as seen by comparing the first and second rows in Figure 7(a). To gain insight into the exacerbated decline in epithelial activity, we further divide the epithelial activity into its components, as seen in the second row of Figure 7(b). One can observe a marked drop in the convective fluxes (from 1.348 nmol.s^-1^.cm^-2^ to 0.58 nmol.s^-1^.cm^-2^), which is due to the notable drop in the tight junction water fluxes (from 16.7 nmol.s^-1^.cm^-2^ to 7.21 nmol.s^-1^.cm^-2^), see Equation (37) and Figure 7(b) (first and second rows). In contrast, there is an intensification in passive activities (from 0.34 nmol.s^-1^.cm^-2^ to 1.02 nmol.s^-1^.cm^-2^, which is due to the changes of the normalised electrical potential differences, which appear in the form of linear and exponential expressions, as described by Equation (39), and seen in Figure 7(c) (first and second rows).

The total electrochemical activity falls from 5.67 nmol.s^-1^.cm^-2^ to 0.59 nmol.s^-1^.cm^-2^, see Figure 7(b). We study the electrochemical activities in the last panel by visualising the relevant components and their changes individually: NHE3, SGLT, and Na^+^-H_2_PO_4_ ^−^. Due to the inhibition of the pumps, there is a significant increase in the sodium cellular concentration (from 19.6 mmol/L to 142.7 mmol/L), which notably decreases the sodium concentration gradient between the lumen (140 mmol/L) and cell, decreasing the electrochemical potential difference of sodium across the apical cell membrane. This, in turn, decreases the activity of the transporters related to the production of sodium fluxes, namely: NHE3 (from 2.05 nmol.s^-1^.cm^-2^ to 0.29 nmol.s^-1^.cm^-2^), SGLT (from 2.05 nmol.s^-1^.cm^-2^ to 0.28 nmol.s^-1^.cm^-2^), and Na^+^-H_2_PO_4_ ^−^ (from 0.11 nmol.s^-1^.cm^-2^ to 0.0143 nmol.s^-1^.cm^-2^).

Here, we highlight a deep insight into the sodium flux variations due to the inhibition of the Na^+^-K^+^ pumps and not other existing solute fluxes. Due to the flexibility of the W-PCT-E model, there is future opportunity for similar analyses to describe the system behavior due to the elimination of other transporters and their impact on the different solutes.

#### iii.2 Inhibition of apical Membrane (MI) Transporters

In this section, we separately eliminate the NHE3 antiporter and apical symporters (SGLT and Na^+^-H_2_PO_4_ ^−^) and then we study the behaviour of the W-PCT-E model by analysing the results for membrane fluxes and cellular concentrations relative to each scenario. In Figure 8, we present the membrane fluxes in the first row and the cellular concentrations in the second row. Figure 8(a) displays the membrane fluxes (ES, IE, IS, ME, MI) and cellular concentrations for the four primary solutes (Na^+^, K^+^, Cl^−^, Glucose) in the case of the original full W-PCT-E model. Figure 8(b) represents the membrane fluxes and cellular concentrations due to the inhibition of the NHE3. In panels (c) and (d), we illustrate the effect of the inhibition of SGLT and Na^+^-H_2_PO_4_ ^−^ transporters on the W-PCT-E model responses, respectively.

**Figure 8.**
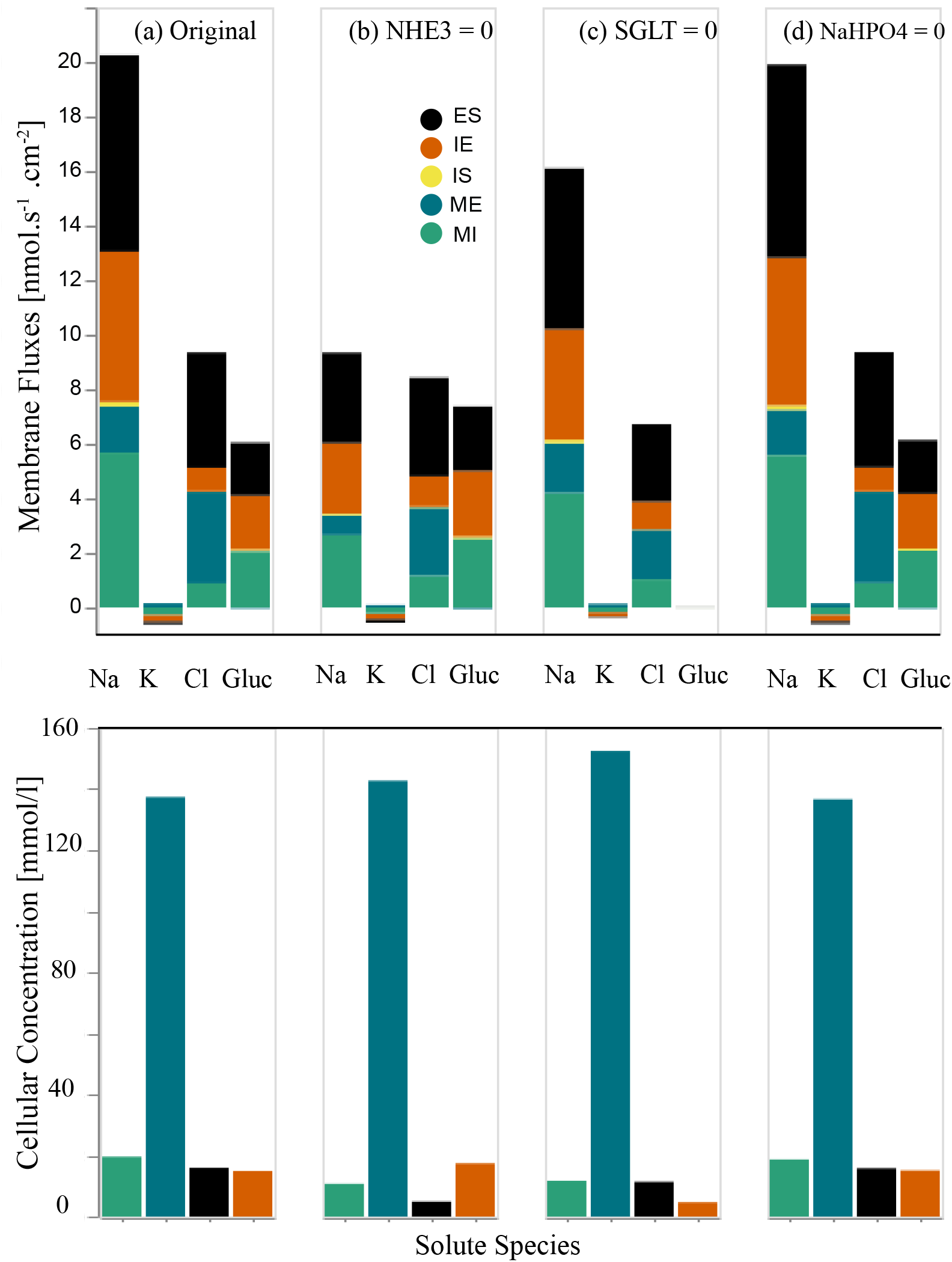
Changes in the membrane fluxes and cellular concentrations due to the inhibition of transporters on the apical cell membrane. In the first row, in each panel, we present four sets of results for four different configurations that depict the total membrane fluxes for the following species: Na^+^, K, Cl, Glucose. The total membrane fluxes include all the membrane activities from five membranes, IS, ME, MI, IE, ES, which are stacked on top of each other. Panel (a) represents the original full model (control scenario). Panel (b) represents flux changes due to the NHE3 elimination. Panel (c) is for the removal of the SGLT, and panel (d) due to inhibition of Na^+^-H_2_PO_4_ ^−^ transporters. In the second row, we illustrate the cellular concentrations for the same species: Na^+^, K^+^, Cl^−^, Glucose in each scenario.

By inhibiting NHE3, one can observe a notable decrease not only in sodium membrane fluxes, but also in cellular sodium and chloride concentrations. While there is an increase in glucose membrane fluxes and cellular glucose concentration, as seen in Figure 8(b).

By eliminating SGLT, we consider the total absence of glucose membrane fluxes which is consistent with the model structure as there are no other sources to produce neither convective nor passive fluxes of glucose. There is a decrease in both Na^+^ and Cl^−^ fluxes; while cellular sodium and chloride concentrations depict decreases, potassium and glucose concentrations demonstrate increases, as can be seen in Figure 8(c).

Removal of Na^+^-H_2_PO_4_ does not show any significant changes neither in epithelial fluxes nor cellular concentrations in any of the four primary solutes, as clearly shown in Figure 8(d).

Here, we focus on the sodium fluxes, as NHE3 is the primary source of sodium fluxes in the epithelial model. NHE3 inhibition stops the exit of hydrogen and ammonia from the cell and the entry of sodium into the cell. Sodium concentration drops from 19.6 mmol/L to 10.5 mmol/L, which confirms the role of NHE3 in the production of Na^+^ fluxes.

To have a better understanding of Figure 8(b) and a deeper insight into the underlying mechanisms of NHE3 and its effect on the sodium fluxes, we study all sources of sodium activity, and the results are featured in the third row in Figure 7.

After the inhibition of NHE3 in Figure 7 (third row), sodium activities in epithelial membranes (ME and MI) drop considerably (the total activity drops from 7.39 nmol.s^-1^.cm^-2^ to 3.37 nmol.s^-1^.cm^-2^, panel 7(a)). To better understand the origin of these changes, we further divide the epithelial sodium activity into convective, passive, and electrochemical component fluxes, as seen in Figure 7(b) (third row).

As we mentioned before, convective and passive fluxes in the epithelial membrane are mainly through the tight junction. In Figure 7(b) (third row), one can observe a notable drop in the convective fluxes (from 1.348 nmol.s^-1^.cm^-2^ to 0.88 nmol.s^-1^.cm^-2^), which is mostly due to the reduction in the water fluxes (from 16.7 nmol.s^-1^.cm^-2^ to nmol.s^-1^.cm^-2^), see Equation (37).

There is a slight reduction in passive fluxes accompanied by changes in direction (from 0.34 nmol.s^-1^.cm^-2^ to−0.15 nmol.s^-1^.cm^-2^). To explain the decrease in the passive fluxes, we need to bring to attention that the main driving force for the passive fluxes is not only the normalised electrical potential differences (which are in the form of linear and exponential components, see Equation (39)) but also solute concentrations. The total electrochemical activity falls from 5.67 nmol.s^-1^.cm^-2^ to 2.64 nmol.s^-1^.cm^-2^, see Figure 7(b). We then investigate the coupled transport fluxes in Figure 7(c) by visualising the components individually: NHE3, SGLT, and Na^+^-H_2_PO_4_ ^−^. Sodium fluxes for NHE3 dropped to zero as a result of inhibition. Due to the changes in sodium cellular concentration (from 19.6 mmol/L to 10.7 mmol/L), the sodium concentration gradient between the lumen (140 mmol/L) and cell increases notably, increasing the electrochemical potential difference of sodium across the apical cell membrane, which increases the activity of transporters related to the production of sodium fluxes (SGLT from 2.05 nmol.s^-1^.cm^-2^ to 2.50 nmol.s^-1^.cm^-2^, and Na^+^-H_2_PO_4_ ^−^ from 0.11 nmol.s^-1^.cm^-2^ to 0.143 nmol.s^-1^.cm^-2^).

## IV. Discussion

Mathematical modelling provides a tool for the investigation of complex physiological phenomena; however, existing mathematical models of epithelial transporters are, in general, not readily findable, accessible, interoperable, nor reusable (FAIR) [20]. The opportunities to reuse these models in future novel studies are therefore remote; thus requiring investigators to first reimplement and/or build their models based on the knowledge that is incomplete, leading to large amounts of resources being required prior to even being able to bring the models to life to address novel applications. For example, to design a specific instance of an epithelial system or even a comprehensive virtual nephron to test different hypotheses or investigate complex diseases.

To help address this, we have presented here what we believe to be a comprehensive and FAIR epithelial model for the PCT of the renal nephron. This model encapsulates and recapitulates the seminal work performed by Weinstein and colleagues [1–5] over many years. In the case of model reproducibility, we have demonstrated that the W-PCT-E model reported here can reproduce many different aspects from related or earlier works. We chose three exemplars [2,3,16], the constant parameters and boundary conditions tuned according to the model of interest. In all cases, we observed a close agreement between the W-PCT-E simulation outcomes and the results reported in previous works, see Section ii (Table 1, Figures 4 and 5).

To show the flexibility of the implementation of the W-PCT-E through the application of structural analysis, we investigated the impact of each transporter on the W-PCT-E model responses (total fluxes and cellular concentrations) through the inhibition of each transporter, see Section iii (Figures 6,7, and 8).

To further demonstrate the comprehensiveness and flexibility of the W-PCT-E model, we now briefly explore various physiological phenomena using our model.

### Clinical studies have shown that excess glucose in the cell and bloodstream is associated with Type II Diabetes (T2D) [21–29]

The inhibition of SGLT for the treatment of T2D [30,31] has shown improvement of glycemic control and T2D [32–34]. According to the W-PCT-E model, the inhibition of the SGLT transporters decreases the cellular Na^+^ and glucose fluxes which decrease the cellular concentration of glucose, see Figure 8(c). We also observed a decrease in interspace concentration of glucose, which would lead to less glucose available for reabsorbtion into the blood. Thus, demonstrating a clear consistency between the W-PCT-E and the clinical findings regarding the inhibition of SGLT and improvement of T2D.

### Clinical reports illustrate that NHE3 residing in the apical membrane mediates transcellular reabsorption of Na^+^ and fluid reabsorption [35,36], and show that NHE3 deficiency can cause a reduction in Na^+^ reabsorption [37,38]

These observations put renal Na^+^ reabsorption via NHE3 in a central position in the development and control of salt loading- and volume expansion-mediated hypertension [39]. We performed some simulation experiments to check that our W-PCT-E model is consistent with these findings by inhibiting NHE3. In doing so, we observed that inhibition of the apical NHE3 decreases the intracellular concentration of HCO_3_ ^−^ and HCO_2_ ^−^, which in turn is transported via the apical Cl^−^ /HCO_3_ ^−^ or Cl^−^ /HCO_2_ ^−^ exchangers. The inhibited apical NHE3 and the Cl^−^ /base exchangers work in parallel and produce net Na^+^ reduction and Cl^−^ reabsorption in the proximal tubule, see Figure 9b)-9(e)((cellular concentrations) and 9(f) (cell volume). These findings are also consistent with other studies demonstrating that the inhibition of either the apical NHE3 and Cl^−^ /base exchangers inhibits net Na^+^ and Cl^−^ absorption in the proximal tubule [40–44].

**Figure 9.**
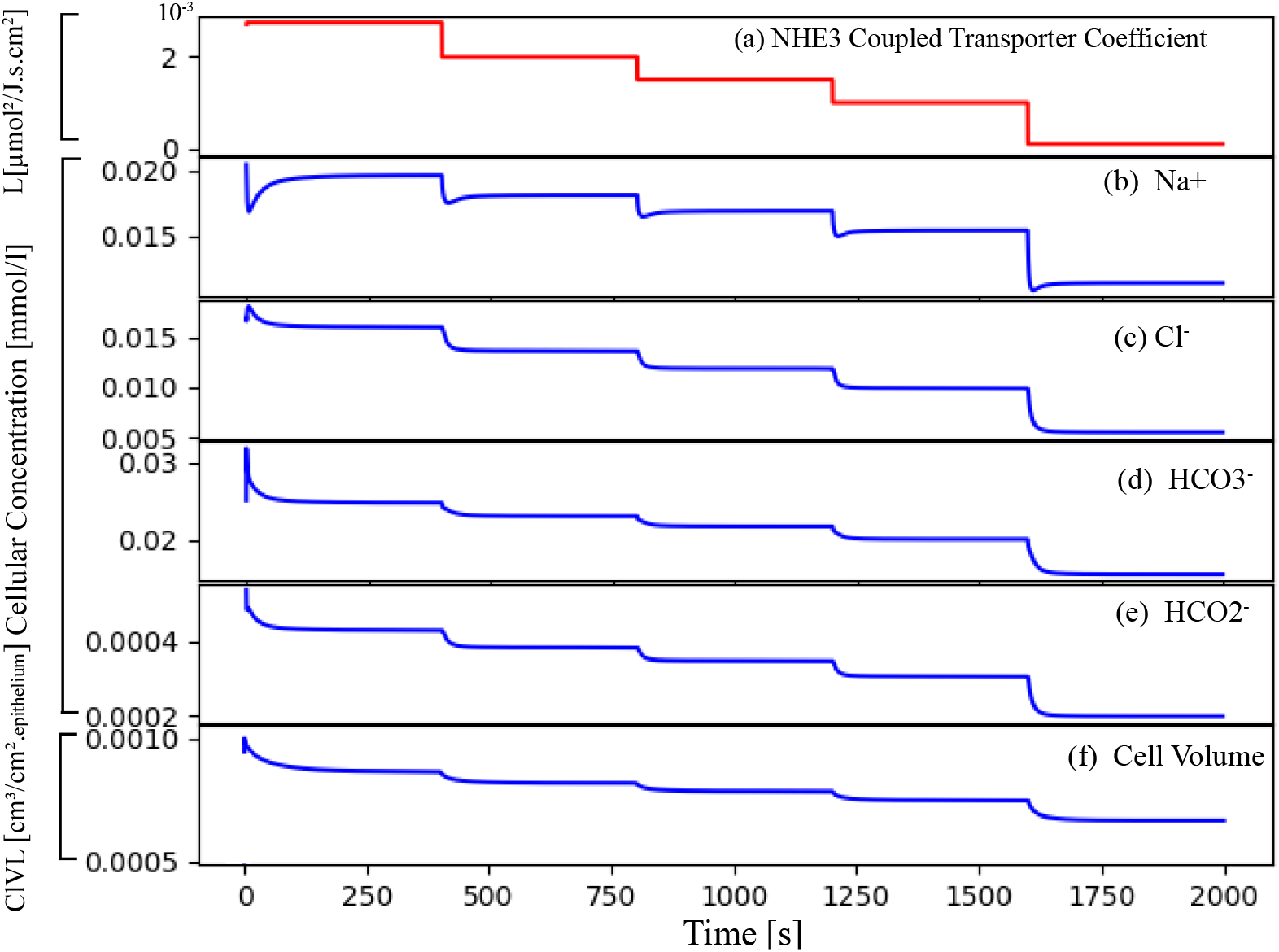
Effect of reduction in NHE3 coupled transporter coefficient on the cellular concentration and cell volumes. Panel (a): NHE3 coupled transporter coefficient decreased in a step-wise manner, at each step there is an decrease of 20% in regard to the original value. Panels (b)-(e): represent changes in some selected cellular solutes concentrations due to the changes in NHE3 coupled transporter coefficient. Panel (f): changes on cell volume.

### Clinical results reveal that a global knockout of NHE3 gene on the proximal tubule reduces water fluxes and HCO_3_ ^−^ absorption [45]

The W-PCT-E model demonstrates that the apical NHE3 exchanger can inhibit HCO_3_ ^−^ in the proximal epithelial model. We designed a set of experiments, in which we decreased the coupled transport coefficient for NHE_3_, see Figure 9. We observed that the inhibition of NHE_3_ lowers the Cl^−^ /HCO_3_ ^−^ and reduces the cell volume. These findings are consistent with the earlier findings [42,45–47].

### Clinical reports show that there is a coordination between basolateral Na^+^/K^+^-ATPase and apical NHE3 activities; they simultaneously regulate the Na^+^transport which can be the cause of inhibition or activation of transepithelial Na^+^ transport [24,39,48–50]

In the W-PCT-E model, one can see that there is strong coordination between basolateral Na^+^/K^+^-ATPase and apical NHE3 activities; inhibition of NHE3 (Na^+^/K^+^-ATPase) can cause the inhibition of Na^+^/K^+^-ATPase (NHE3). We can conclude from this observation that in the W-PCT-E model, sodium is primarily reabsorbed via NHE3 which is regulated by Na^+^/K^+^-ATPase, see supplementary information Figure 10.

**Figure 10.**
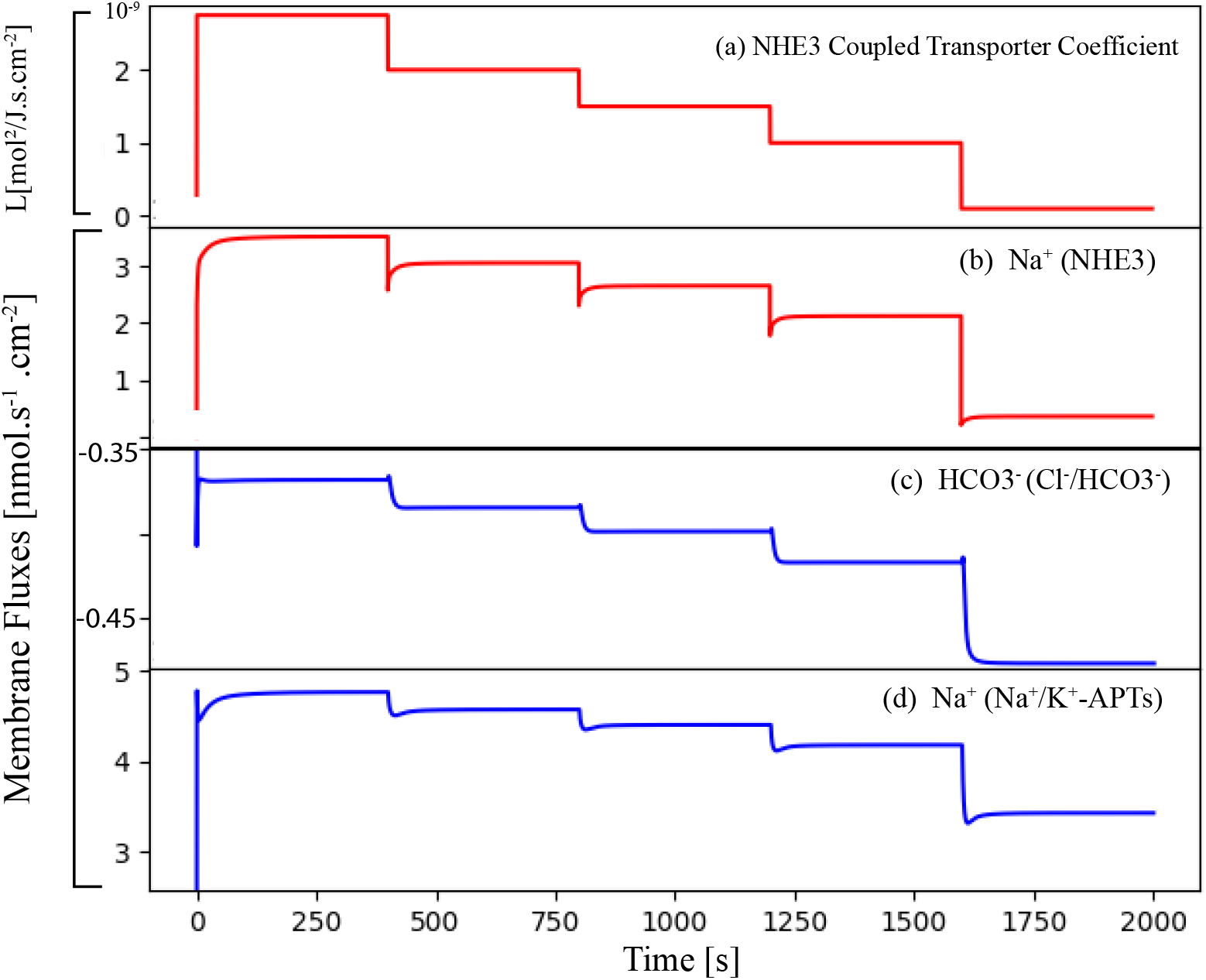
Effect of reduction in NHE3 coupled transporter coefficient on the membrane fluxes. Panel (a): NHE3 coupled transporter coefficient decreased in a step-wise manner, at each step there is an decrease of 20% in regard to the original value. Panels (b)-(d): represent changes in some selected membrane fluxes for selected transporters due to the changes in NHE3 coupled transporter coefficient.

### The kidney plays a critical role in the regulation of body electrolyte and fluid balance, primarily occurring in the proximal tubule segment of the nephron [51–53]

A balance of body extracellular electrolyte composition and fluid volume is essential for all animals and humans to survive. Either excess or shortage of crucial extracellular electrolytes or overall fluid volume may lead to disturbance of blood pressure and abnormalities in cellular functions, including cell volume [53–57]. The W-PCT-E model can capture the impact of modulation of different transporters on the fluid balance and cellular volume. For example, with the inhibition of basal-cell Na^+^/K^+^-ATPase or the activation of apical NHE3 or SGLT, we observed growth in the cell volume. Such observations are consistent with previous modelling approaches and clinical reports [24,55–58]. Building on the capability of the present W-PCT-E model to predict such volume changes, we believe this is an area with great potential to explore in our future work concerning obesity and hypertension [54,59,60].

The implementation of the present epithelial transport model to support the composition and parameterisation of the proximal tubule epithelial system (the W-PCT-E model) is available on Github under a license which allows open and unrestricted reuse. The implementation is in Python (version 3) to make it broadly accessible. As demonstrated above, the implementation is sufficiently flexible and configurable to support the generation of different epithelial models. Users need only to extract the required constituent transporter modules from the available repository and then integrate them into the desired configurations. The boundary conditions need to be changed according to the epithelium of interest. In sharing the model implementation between the authors of this manuscript, we tested for reusability as we each work with W-PCT-E model.

## V. Conclusions

We believe the time is now right to develop a reproducible and FAIR virtual nephron Here we term this the iNephron. It is critical to ensure that the capabilities of published models are captured while being sufficiently flexible and configurable to support the generation of novel models to investigate specific scientific or clinical questions. In achieving this, iNephron will provide a tool to investigate the fundamental mechanisms involved in hypertension, diabetes, and many other kidney diseases. Furthermore, iNephron will be developed and shared following established best practices in open and reproducible science to guarantee that the scientific community can benefit and extend from this work to improve our collective understanding of these diseases.

The work we presented here is our first step toward achieving the iNephron. As we look to grow the repository of available transporter modules, we also look to better follow FAIR principles [20,61], and move toward a standards-based repository of reusable modules (e.g., [62]). To ensure the composed models are meaningful and that model composition can occur reliably, we are planning to migrate our repository of reusable modules to ensure thermodynamic consistency [8–12]. This is a requirement for the arbitrary composition of models needed for iNephron.

As we have shown, the W-PCT-E model is actually a generic epithelial model which is flexible and configurable to support the generation of different epithelial models, where the user is able to meet their design requirements, easing the process for “getting started” with a novel modelling study. All one needs is to provide the model with a set of transporters and boundary conditions appropriate for the epithelial model of interest. Additionally, by establishing a comprehensive ability to perform sensitivity analyses, we provided tools by which future users are able to test their own additions or modifications of this model with confidence. Our testing, summarised in this manuscript with more detailed findings in the supplemental material, shows that our model behaves as expected in physiological terms.

## Availability

The W-PCT-E Python code can be found here: https://github.com/iNephron/W-PCT-E. In that Github repository, we also provide documentation explaning how to reproduce the results presented here and suggesting how the model may be reused. The results presented here specifically use Release of this implementation, available from: https://github.com/iNephron/W-PCT-E/releases/tag/v1.0.0.

## Supporting information

## Acknowledgments

PJB acknowledges the support of the Brazilian agencies CNPq (grant numbers 301224/2016-1 and 407751/2018-1), and FAPESP (grant numbers 2014/50889-7 and 2018/14221-2). SS acknowledges the financial support provided by the Aotearoa Foundation.

## Footnotes

1 The membrane charge constraint will be added to the system through equation (35), which elevates the total number of equations to 35.

